# Topoisomerase IIb binding underlies frequently mutated elements in cancer genomes

**DOI:** 10.1101/2024.11.06.622320

**Authors:** Liis Uusküla-Reimand, Christian A. Lee, Robin H. Oh, Zoe P. Klein, Nina Adler, Sana Akhtar Alvi, Ellen Langille, Elisa Pasini, Kevin C. L. Cheng, Diala Abd-Rabbo, Huayun Hou, Ricky Tsai, Mamatha Bhat, Daniel Schramek, Michael D. Wilson, Jüri Reimand

**Author notes:** These authors contributed equally (Liis Uusküla-Reimand, Christian A. Lee).

## Abstract

Type-II topoisomerases resolve topological stress in DNA through controlled double-strand breaks. While TOP2A is a chemotherapy target in proliferating cells, the ubiquitously expressed TOP2B is a potential off-target. Here we explore roles of TOP2B in mutagenesis by generating DNA-binding maps of TOP2B, CTCF, and RAD21 in human cancer samples and analysing these maps for driver mutations and mutational processes in 6500 whole cancer genomes. TOP2B-CTCF-RAD21 and TOP2B-RAD21 sites are enriched in somatic mutations and structural variants (SVs), especially at evolutionary conserved sites displaying high transcription and long-range chromatin interactions. TOP2B binding underlies SVs and hotspot mutations in cancer-driving genes such as *TP53*, *MYC*, *FOXA1*, and *VHL*, and many cis-regulatory elements. We show that the TOP2B-bound mutational hotspot at *RMRP* drives tumor initiation and growth *in vivo*. These data highlight TOP2B as a protector of the genome from topological challenges whose aberrant activity promotes driver and passenger mutations in cancer genomes.

## INTRODUCTION

Chromatin interactions during transcription, replication, and other cellular processes impose mechanical stress on DNA, leading to entanglements and knots that cause genome instability and cell death if left unresolved. Topological constraints in mammalian genomes are resolved by type II topoisomerases TOP2A and TOP2B that catalyse reversible DNA DSBs and strand passage of uncleaved DNA duplexes ^1^. TOP2A is expressed in proliferating cells and ensures appropriate chromosome condensation and segregation during mitosis. In contrast, TOP2B is expressed ubiquitously across tissues and cell types ^2^. Notably, TOP2B binds active promoters, enhancers, and boundaries of topologically associating domains, often in association with the insulator protein CTCF and the cohesin complex subunit RAD21 ^3,4^, linking TOP2B functions with tissue-specific gene regulation and chromatin architecture ^5–8^. During its reaction cycle, TOP2 temporarily attaches to the cleaved ends of DNA to form a covalent TOP2-DNA cleavage complex ^9^. Inhibiting the reversal of the complex in proliferating cells by anthracycline class of chemotherapy drugs such as doxorubicin and epirubicin results in cytotoxic DSBs and apoptosis in solid and hematological cancers ^10,11^. However, off-target stabilisation of the TOP2B cleavage complexes in non-dividing cells can result in structural variants (SVs) due to aberrant genesis and repair of DSBs, such as the *MLL* (*KMT2A*) translocations commonly found in therapy-induced leukemias ^12–14^. Emerging evidence indicates that TOP2B-mediated DNA lesions and translocations can also originate from endogenous cellular processes such as RNA Polymerase II (Pol II) pause release and transcriptional elongation ^15–20^, as seen in prostate and breast cancers ^21,22^. Despite this evidence, the roles of TOP2B in cancer mutagenesis remain largely unexplored.

Cancer is driven by somatic mutations such as single nucleotide variants (SNVs), insertions-deletions (indels), as well as SVs of copy number alterations (CNAs), inversions, and translocations. Driver mutations deregulate key cancer-related pathways by aberrantly activating oncogenes or disrupting tumor suppressors and can be identified in cancer genomics datasets through elevated mutation rates indicative of positive selection ^23–25^. Most known driver mutations affect protein-coding sequences, whereas fewer non-coding driver mutations have been identified to date ^24–29^. Cancer genomes typically contain a handful of driver mutations, whereas most mutations are considered functionally neutral passengers that result from diverse mutational processes. These mutational processes affect the genome at multiple levels, including nucleotide-level mutational signatures ^30^. They can also result in local mutational enrichments, often observed in regulatory elements such as CTCF binding sites ^31,32^. Furthermore, these can lead to megabase-level variations in mutation burden that correlate with chromatin accessibility and DNA replication in a tissue-specific manner ^33–36^. Mutational processes have been linked to aging, DNA repair deficiencies, carcinogen exposures, or chemotherapies ^37–40^. However, many of these processes remain poorly understood in terms of their underlying mechanisms and etiology.

Here, we study the role of TOP2B in mutational processes and its potential impact on cancer driver mutations across thousands of cancer genomes. Leveraging a comprehensive genome-wide map of TOP2B binding obtained from human tumor biopsies, we reveal a striking connection between TOP2B binding sites and genomic enrichment in somatic mutations and SVs, particularly at conserved, constitutively active regulatory elements marked by CTCF or cohesin binding. Within this landscape, we identify hundreds of individual sites as significant mutational hotspots, often affecting cancer driver genes. We validate one mutational hotspot as a novel cancer driver *in vivo*. These findings shed light on the mutational processes involving TOP2B, potentially explaining how tissue-specific and chemotherapy-mediated DNA damage can lead to mutations at critical regulatory and architectural elements.

## RESULTS

### Conserved DNA-binding landscape of TOP2B in human cancer cells

We generated genome-wide DNA-binding profiles of TOP2B, CTCF, and RAD21 in human liver hepatocellular carcinoma (HCC) samples using chromatin immunoprecipitation sequencing (ChIP-seq). Peak calling across the biological replicates and merging the peaks by factor binding revealed 72,351 unique binding sites (FDR < 0.05; FC ≥ 2). Collectively, we refer to all these sites as TOP2B or CTCF or RAD21 binding sites (TCRBS). 61% of sites were bound by TOP2B including 16,534 *‘triple sites’* bound by TOP2B-CTCF-RAD21 (23%), 19,265 *‘double sites’* bound by TOP2B-RAD21 (27%), and 8,026 sites bound by TOP2B alone (11%) (**Figure 1a**). We also identified 10,609 sites bound by CTCF-RAD21 and 17,054 sites bound by RAD21 alone. Frequent DNA-binding interactions of TOP2B, CTCF, and RAD21 are consistent with our previous study in mouse liver ^3^.

**Figure 1.**
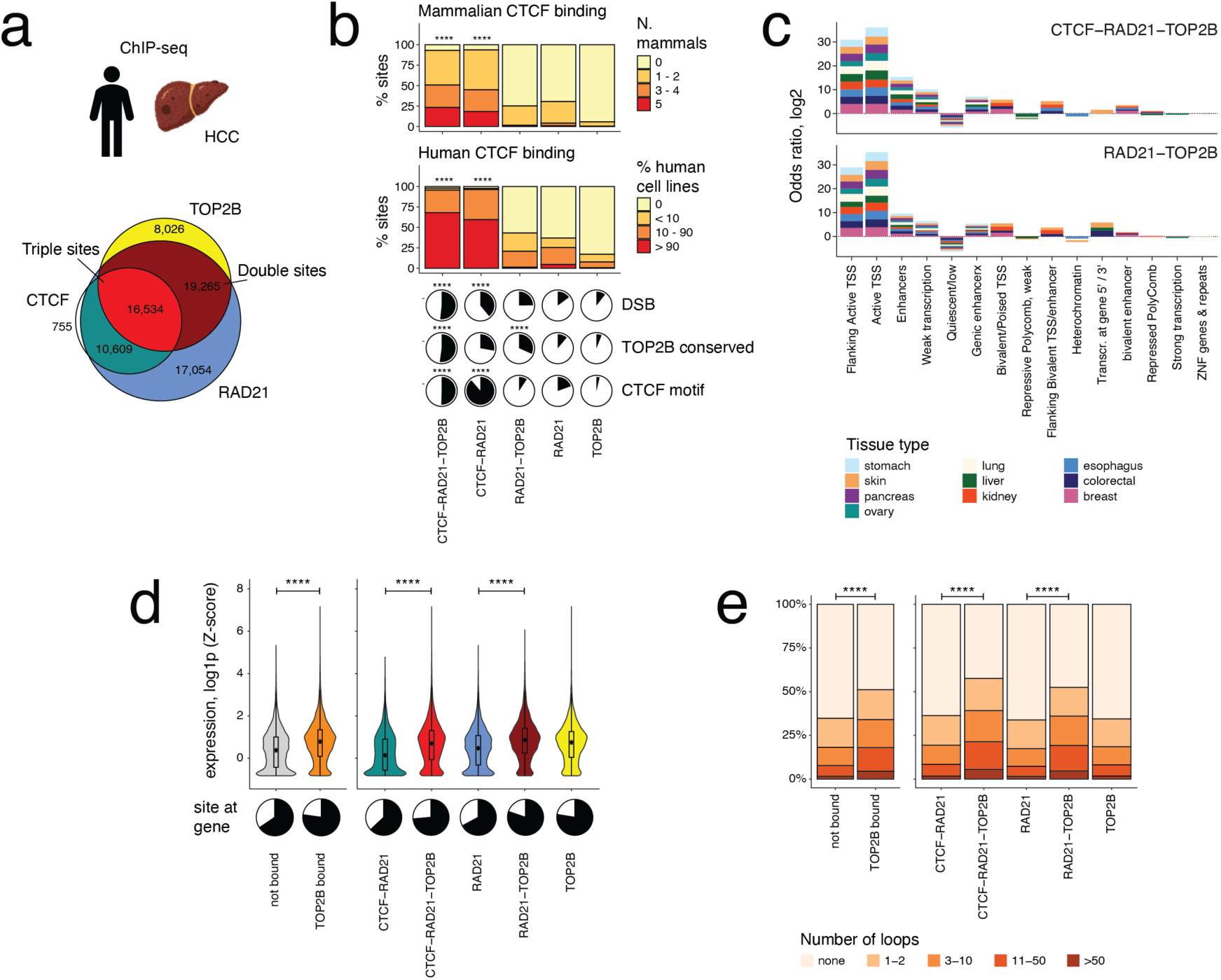
Genome-wide binding landscape of TOP2B in human cancer cells. **(a)** Overview of TOP2B, CTCF, and RAD21 binding sites (TCRBS) identified in clinical samples of human hepatocellular carcinoma using ChIP-seq. The Venn diagram shows TCRBS co-bound by TOP2B-CTCF-RAD21 (*i.e.*, triple sites), TOP2B-RAD21 (*i.e.*, double sites), or in other combinations. **(b)** Evolutionary conservation, pan-tissue activity, and functional characteristics of TCRBS. Triple sites and CTCF-RAD21 sites are conserved based on DNA-binding profiles of CTCF and TOP2B in mammalian livers, occur frequently at constitutively active CTCF binding sites measured from 70 human cell lines, and are enriched in DSB sites in human cell lines. **(c)** TOP2B binding is enriched in active regulatory elements in multiple tissue types. Epigenetic annotations of triple sites and double sites were compared to sites lacking TOP2B binding (i.e., triple *vs* CTCF-RAD21 sites, double *vs* RAD21 sites, respectively). Odds ratios of significantly enriched epigenetic states are shown (hypergeometric test, FDR < 0.05). **(d)** TOP2B binding associates with higher gene expression in cancer. Violin plots show mean gene expression levels at TCRBS based on pan-cancer transcriptomics datasets from PCAWG. Pie charts below show the fractions of sites at genes. **(e)** TOP2B binding associates with more frequent promoter-enhancer chromatin interactions across 27 human cell types. P-values of Wilcoxon rank-sum tests are shown in panels (d-e).

To gain insights into the functional properties of TCRBS, we first studied their evolutionary conservation and epigenetic activity using CTCF binding maps in 70 human cell lines in ENCODE ^41^ and evolutionary maps of CTCF binding in mammalian livers ^42^. Triple sites showed highly significant levels of evolutionary conservation and activity across human cell types (**Figure 1b**): 71% of these occurred at constitutively active CTCF binding sites found in 90% of human cell lines, and 35% showed conserved CTCF binding in four or more mammalian livers. Also, 50% of triple sites had a CTCF DNA-binding motif, 52% showed conserved TOP2B binding in mouse liver ^3^, and 51% occurred at DSB sites from earlier cell line experiments ^4^, collectively underscoring the functional importance of triple sites in cells and their targeting by topoisomerases. CTCF-RAD21 sites were also conserved and enriched in DSBs, potentially because these also include TOP2B binding sites that remain undetected due to technical limitations. In contrast to triple sites, double sites and TOP2B-only sites overlapped with fewer evolutionarily conserved or constitutively active CTCF binding sites, which is consistent with no CTCF binding at these sites in our data. However, double sites (32%) were also significantly enriched in sites with conserved TOP2B binding in mouse liver, suggesting that CTCF binding alone does not fully recapitulate TOP2B binding. CTCF-independent binding of cohesin at cis-regulatory elements is involved in tissue-specific transcription regulation ^43,44^, and double sites may therefore represent a different mode of TOP2B activity mediated by cohesin in tissue-specific loci.

We characterised the chromatin states of TCRBS in ten major tissue types from the Roadmap Epigenomics Project ^45^. To exclude potential biases of CTCF or RAD21 binding, we examined TOP2B-bound sites relative to control sites lacking TOP2B binding, comparing triple sites to CTCF-RAD21 sites and double sites to RAD21-only sites. Both triple sites and double sites were highly enriched in active transcription start sites (TSS) and their flanking regions in all tissue types (FDR < 0.05) (**Figure 1c**). For example, 27% of triple sites and 37% of double sites occurred at active TSSs in liver (17% and 23% sites expected, respectively; hypergeometric P < 2.2 x 10^-16^). In contrast, quiescent, heterochromatin, and repressive states were depleted at TOP2B binding sites. Although our TOP2B binding maps were obtained from liver cancer samples, regulatory chromatin states of other tissue types were also enriched, indicating that TOP2B often binds to *cis*-regulatory elements shared across tissues and cell types.

We examined transcriptional and gene-regulatory attributes of TOP2B binding. First, we studied pan-cancer gene expression data from 2,851 transcriptomes of primary and metastatic cancers from Pan-cancer Analysis of Whole Genomes (PCAWG) and Hartwig Medical Foundation (HMF) projects ^46,47^. TOP2B binding sites occurred more often adjacent to genes compared to sites lacking TOP2B binding (77% *vs*. 65%, hypergeometric P < 2.2 x 10^-16^) and mean gene expression at TOP2B binding sites was significantly higher (Wilcoxon test, P < 2.2 x 10^-16^) (**Figure 1d**). Similarly, gene expression at triple sites and double sites was higher compared to control sites lacking TOP2B binding. TOP2B binding also associated with higher gene expression in individual cancer types and in primary and metastatic cancers (**Figure S1**).

Next, to investigate TOP2B binding in the context of chromatin architecture, we studied promoter-enhancer interactions of 27 normal tissue types from a promoter-capture HiC dataset ^48^. TOP2B binding sites occurred at hotspots of regulatory chromatin interactions: 51% of sites had at least one promoter-enhancer interaction and 18% of sites had ten or more interactions, significantly exceeding sites not bound by TOP2B (35% and 8%, respectively, Wilcoxon P < 2.2 x 10^-16^) (**Figure 1e**). We compared triple sites and double sites to control sites lacking TOP2B binding and again found that TOP2B sites had more frequent promoter-enhancer interactions, extending our prior findings in mouse liver ^3^. Thus, TOP2B binding in human cancer cells occurs at transcriptionally active sites with three-dimensional chromatin interactions that are bound by CTCF in many human tissue types, show conserved TOP2B or CTCF binding in mammalian epigenomes, and coincide with *in vitro* DSB maps attributed to topoisomerase activity, suggesting that TOP2B may alter functional and structural genomic elements in cancer genomes.

### TOP2B binding sites are enriched in SNVs and indels in cancer genomes

To understand the role of TOP2B in cancer mutagenesis, we studied somatic mutations of 6,495 cancer samples of 18 major cancer sites profiled by whole-genome sequencing (WGS), including 2,452 primary cancers from the PCAWG project ^26^ and 4,043 metastatic cancers from the HMF project ^47^, with 97 million SNVs and indels in total. First, we characterised local mutational processes of small mutations in TCRBS using the computational method RM2 ^32^, which evaluates mutational enrichments across genomic sites relative to flanking control sequences using trinucleotide and megabase-level covariates (**Figure 2a**). Triple sites were consistently most enriched in mutations, as observed in cancers of the liver, esophagus, lung, colon, breast, prostate, skin melanomas, and pan-cancer (FDR < 0.05) (**Figure 2b**). Double sites and CTCF-RAD21 sites were also enriched in mutations although at lower significance levels. In contrast, sites only bound by RAD21 or TOP2B generally showed no mutational enrichments, while too few CTCF-only sites were available for this analysis. Site types were characterised by distinct patterns of localised mutagenesis: triple sites and CTCF-RAD21 sites had focal mutation enrichments within hundred base pairs of peak summits, while double sites had a smoother and wider increase in mutation burden (**Figure 2c**).

**Figure 2.**
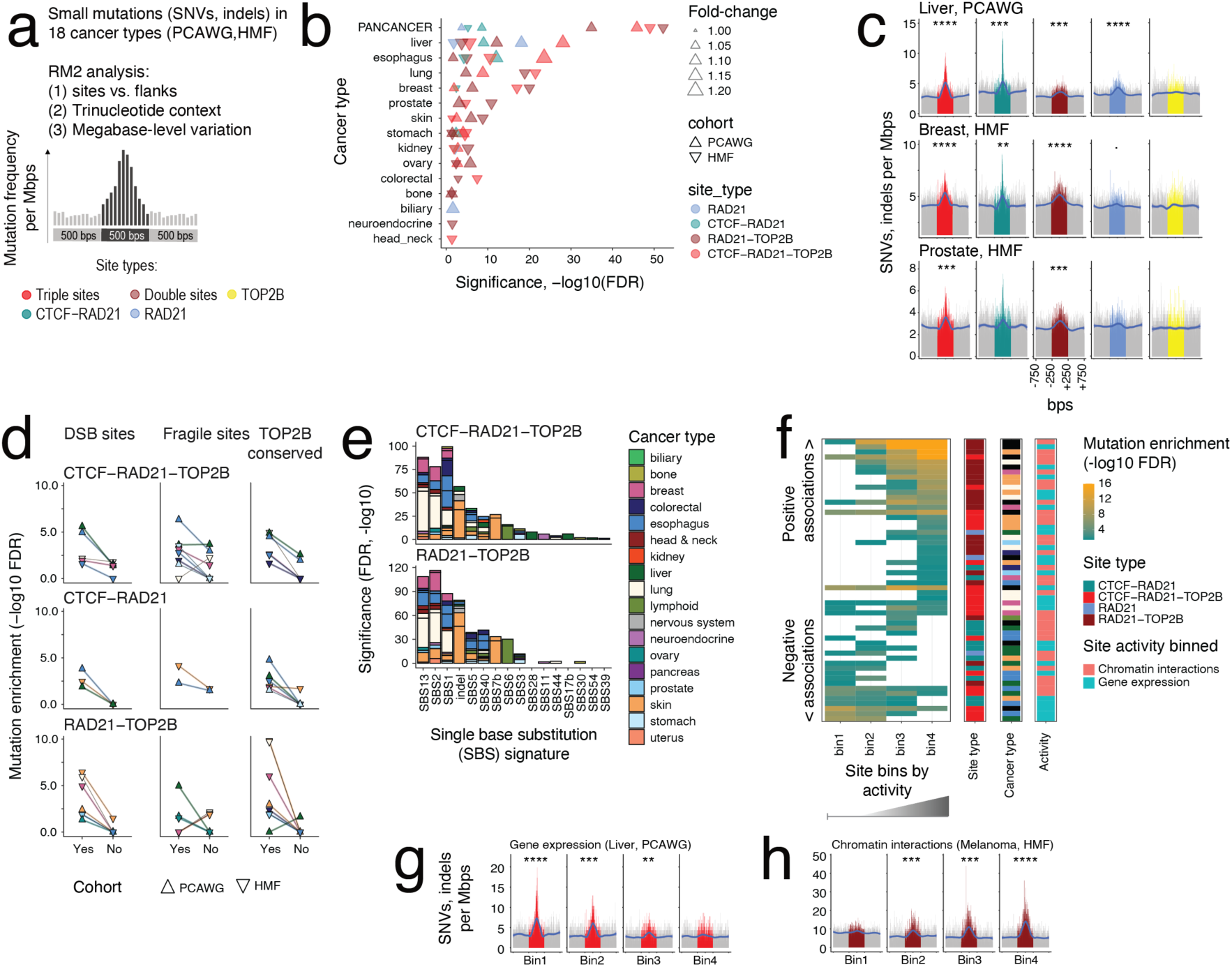
Mutational processes of SNVs and indels at TCRBS in cancer genomes. **(a)** Detecting mutational processes at TCRBS. Binding sites were aligned and compared to flanking control sequences using the RM2 method, which quantifies mutational enrichments across sites using their trinucleotide context and megabase-scale mutation burden. **(b)** Enrichments of SNVs and indels in TCRBS in primary and metastatic cancers (RM2, FDR < 0.001). **(c)** Examples of mutational enrichments in primary liver cancers (top), metastatic breast cancers (middle), and metastatic prostate cancers (bottom). Grassy-hills plots show mutation burden across all sites of 250-bps length (colored area) relative to flanking sequences ±250 bps around the sites (grey area). **(d)** Functional annotations of TOP2B binding sites associate with increased mutagenesis. Sites with and without functional annotations are compared: double-strand break (DSB) sites (left), common fragile sites (middle), and conserved TOP2B sites in mice (right). The Y-axis shows significance of local mutation rate increase at sites relative to control regions flanking the sites (RM2, FDR < 0.05). **(e)** Mutational signatures of single base substitutions (SBS) are enriched in TOP2B binding sites. Triple sites are compared to CTCF-RAD21 sites and double sites are compared to RAD21 sites and significant enrichments are reported (FDR < 0.05). **(f)** Epigenomic and transcriptomic activity at TOP2B binding sites correlates with somatic mutagenesis. Sites were grouped by transcriptional activity or chromatin interaction frequency into four bins (*i.e.*, none, low, middle, high) and local mutagenesis in each bin was evaluated using RM2. The heatmap shows site categories sorted by increasing or decreasing mutagenesis with respect to transcription or chromatin interactions (top vs. bottom). **(g-h)** Examples of mutational processes in TOP2B binding sites associating with transcription and chromatin interactions. **(g)** In liver cancer, triple sites with low or no transcription have higher rates of local mutagenesis while sites at highly expressed genes have no mutational enrichments at sites. **(h)** In melanoma, double sites with high frequency of chromatin interactions are strongly enriched in mutations while sites with few or no chromatin interactions show no mutational enrichments. P-values from RM2 are shown.

We examined the functional annotations of TOP2B binding sites in the context of cancer mutations. Higher levels of somatic mutagenesis were observed at triple sites and double sites associated with *in vitro* DSB activity ^4^, genomic fragile sites ^49^, and conserved TOP2B binding in mouse liver ^3^, compared to triple sites and double sites lacking these functional annotations.

Thus, such functional characteristics define subsets of TOP2B binding sites with even higher local mutagenesis (**Figure 2d**). Similar albeit weaker trends were observed at CTCF-RAD21 binding sites. Thus, TOP2B binding may explain the increased mutagenesis at CTCF and cohesin binding sites observed previously in many cancer types ^31,32^.

Specific mutational signatures of single base substitutions (SBS) were also enriched in TOP2B binding sites relative to control combinations of sites lacking TOP2B binding (**Figure 2e**).

APOBEC signatures SBS2 and SBS13 showed the strongest enrichments and were especially prominent in lung, esophageal and breast cancers, suggesting a link between TOP2B binding, advanced cancers ^50^ and therapy-associated mutagenesis by APOBEC enzymes ^51^. UV light signatures and indel mutations in TOP2B-bound sites were enriched in melanomas, and clock-like signatures SBS1, SBS5 and SBS40 were also found enriched at TOP2B binding sites in multiple cancer types.

We asked if the mutational processes at TOP2B binding sites associated with their transcriptional or chromatin context. We leveraged 2,851 matched cancer transcriptomes from PCAWG and HMF and grouped all TCRBS into four bins based on either silent, low, medium, or high gene expression in specific cancer types. Mutagenesis at TOP2B binding sites was associated with both higher and lower gene expression levels depending on cancer types (**Figure 2f**). In liver and esophageal cancers, triple sites in transcriptionally silent or lowly expressed loci were highly enriched in somatic mutations while sites at highly expressed genes were not (**Figure 2g, Figure S2a**). In contrast, an opposite pattern was observed in melanoma where double sites at highly expressed genes were strongly enriched in mutations while transcriptionally silent sites were not enriched (**Figure S2b**). Similarly, context-specific interactions with long-range chromatin interactions were apparent at TOP2B binding sites (**Figure 2f**). We grouped TCRBS based on chromatin interactions identified from 27 human cell types ^48^ and compared sites with four levels of chromatin interaction activity. Similarly to gene expression, positive and negative associations with mutagenesis were seen. For example, double sites with many chromatin interactions were highly enriched in mutations while sites with no or few interactions were not enriched, as was seen in skin and breast cancer (**Figure 2h**, **Figure S2d**). In contrast, esophageal and other digestive tract cancers often showed opposite associations where the TOP2B binding sites with few or no chromatin interactions were most enriched in mutations (**Figure S2c**).

Collectively, these data show that TOP2B-bound triple sites and double sites are highly enriched in somatic SNVs and indels in multiple cancer types. Conserved TOP2B binding and genomic DSBs at the sites provide evidence of topoisomerase activity ^4,5,7,21,52^, while the transcriptional and chromatin associations of mutational processes may be determined by context-specific mutagenesis, such as transcription-coupled nucleotide excision repair that is modulated by transcriptional activity, RNA Pol II pause-release^18,21^, or transcription factor (TF) binding ^53,54^.

### Structural variants are enriched at TOP2B binding sites

We studied TCRBS in the context of 2.1 million structural variant (SV) breakpoints (SVBPs) in primary cancers and metastases. TOP2B binding sites had significantly more pan-cancer SVBPs compared to genome-wide tiling windows, and triple sites and double sites also had significantly more SVBPs than sites lacking TOP2B binding (P < 2.2 x 10^-16^, Wilcoxon test) (**Figure 3a**). We confirmed these findings using more stringent analyses in each cancer type in which we compared the distribution of SVBPs in TOP2B binding sites relative to the high burden of SNVs and indels we observed above. Again, triple sites and double sites were enriched in SVBPs compared to control sites lacking TOP2B binding (hypergeometric test, FDR < 0.05) (**Figure 3b**). SVBP enrichment was apparent in all 18 cancer types and primary and metastatic cohorts, especially in esophageal, lung, colorectal, skin, and prostate cancers. Thus, TOP2B binding sites are enriched in SVBPs beyond what is explained by the binding of CTCF or RAD21.

**Figure 3.**
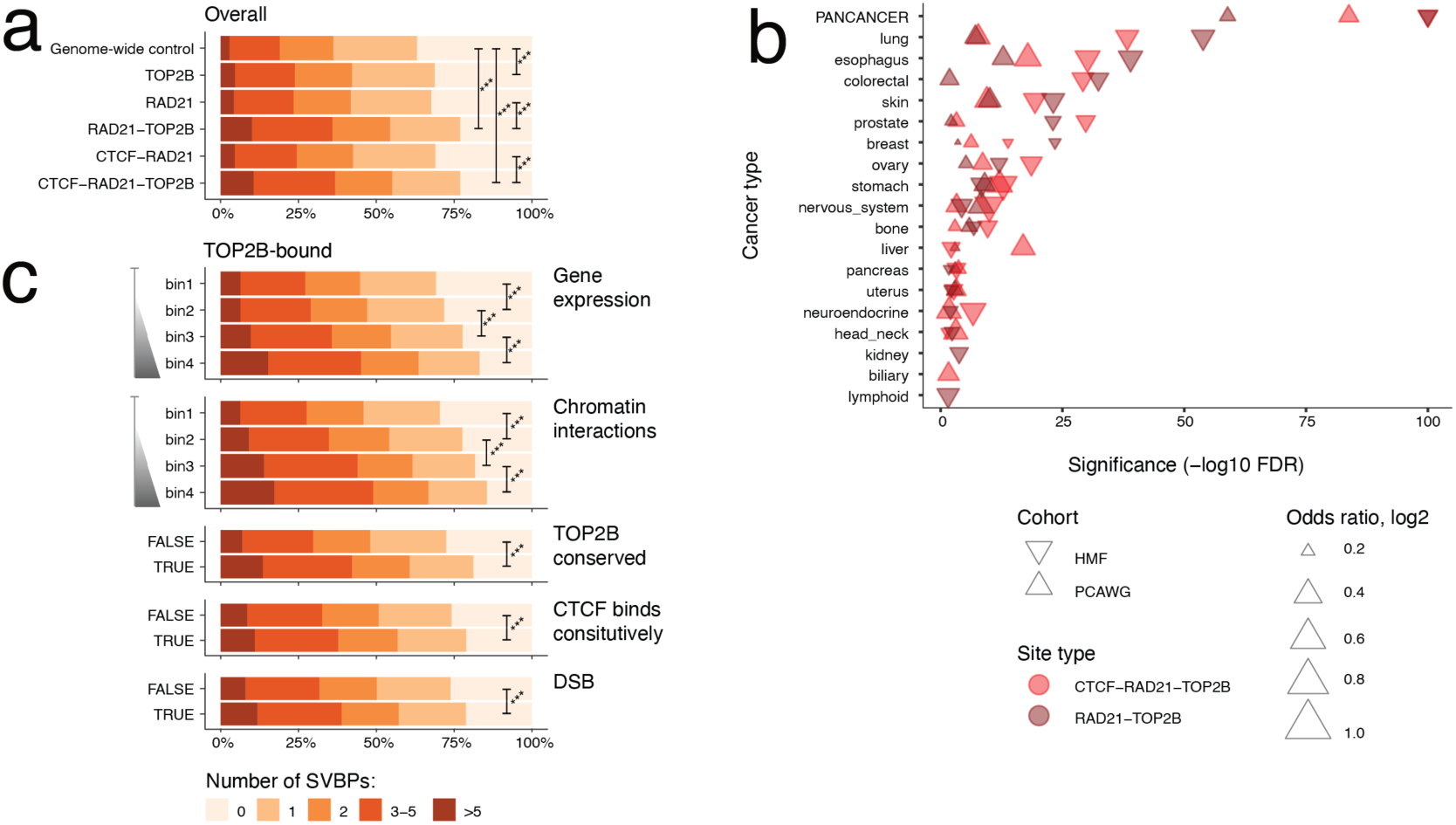
Structural variant (SV) breakpoints (SVBPs) are enriched at TOP2B binding sites in cancer genomes. **(a)** Comparison of SVBP frequency in TOP2B binding sites in pooled pan-cancer data from primary cancers and metastases. The stacked bar plots show the fractions of sites with high, medium, or low number of SVBPs. Focal SVBP burden at sites was compared in TOP2B-bound sites and non-bound sites (triple sites vs CTCF-RAD21 sites, double sites vs RAD21 sites). As genome-wide controls, tiled genomic windows of matching size were used. Wilcoxon rank-sum P-values are shown. **(b)** Enrichment of SVBPs in triple sites and double sites relative to SNV/indel distributions in the sites (hypergeometric test, FDR < 0.05). Triple and double sites are compared to sites lacking TOP2B binding (*i.e.*, triple sites *vs.* CTCF-RAD21 sites; double sites *vs.* RAD21 sites). **(c)** SVBPs in TOP2B binding sites are more frequent in functionally active sites. Sites were grouped into four bins by pan-cancer gene expression and frequency of chromatin interactions shown in top two facets as well as evolutionary conservation, constitutive CTC F binding, and double strand break (DSB) activity. Sites lacking these annotations were used as controls. P-values from Wilcoxon tests are shown.

We focused on TOP2B binding sites and asked if their functional annotations were informative of SVBP burden. Sites at highly expressed genes had significantly more pan-cancer SVBPs compared to sites at lowly expressed genes or intergenic sites (Wilcoxon test, P < 2.2 x 10^-16^), results that were recapitulated in most cancer types (**Figure 3c, Figure S3**). Similarly, TOP2B binding sites at sites with frequent promoter-enhancer interactions were enriched in SVBPs compared to sites with fewer or no interactions, and TOP2B binding sites with *in vitro* DSB activity ^4^ or conserved TOP2B or CTCF binding in mice ^3^ were also enriched in SVBPs (P < 2.2 x 10^-16^). Thus, TOP2B binding sites at evolutionarily conserved, highly transcribed, or frequently interacting chromatin elements are hotspots of genomic rearrangements, potentially due to topoisomerase activity that resolves topological constraints through DSB genesis and re-ligation.

### Recurrently mutated TOP2B binding sites in cancer

Somatic driver mutations undergo positive selection in cancer genomes and driver genes are characterised by mutation rates that significantly exceed expected background mutations. We asked if individual TOP2B binding sites were significantly enriched in somatic mutations by performing driver analyses of individual cancer types in the PCAWG and HMF datasets. Cancer driver analysis using the ActiveDriverWGS method ^25^ revealed 243 unique sites from our catalogue of TOP2B, CTCF or RAD21 binding sites that we refer to as frequently mutated regulatory elements (FMREs). These included 189 FMREs with enriched SNVs or indels, and 65 FMREs with enriched SVBPs, and a few FMREs found in both analyses (ActiveDriverWGS FDR < 0.05) (**Figure 4a**). FMREs were detected in 17 of 18 cancer types we studied, and most were identified in skin melanomas, and cancers of colon, breast, esophageal, liver, lymph, and prostate. FMREs were enriched in known driver genes (42 observed *vs*. 8.6 expected; hypergeometric P = 2.9 x 10^-19^). These included major pan-cancer drivers such as *TP53*, *MYC*, *PTEN* and *EGFR* as well as tissue-specific drivers such as *VHL*, *TMPRSS2*, and *FOXA1*. The FMREs included SNVs and indels with protein-coding and non-coding impact as well as frequent SV breakpoints. SV-based FMREs showed a high rate of translocations that collectively affected 31 known cancer genes in 404 cancer samples (**Figure 4e**).

**Figure 4.**
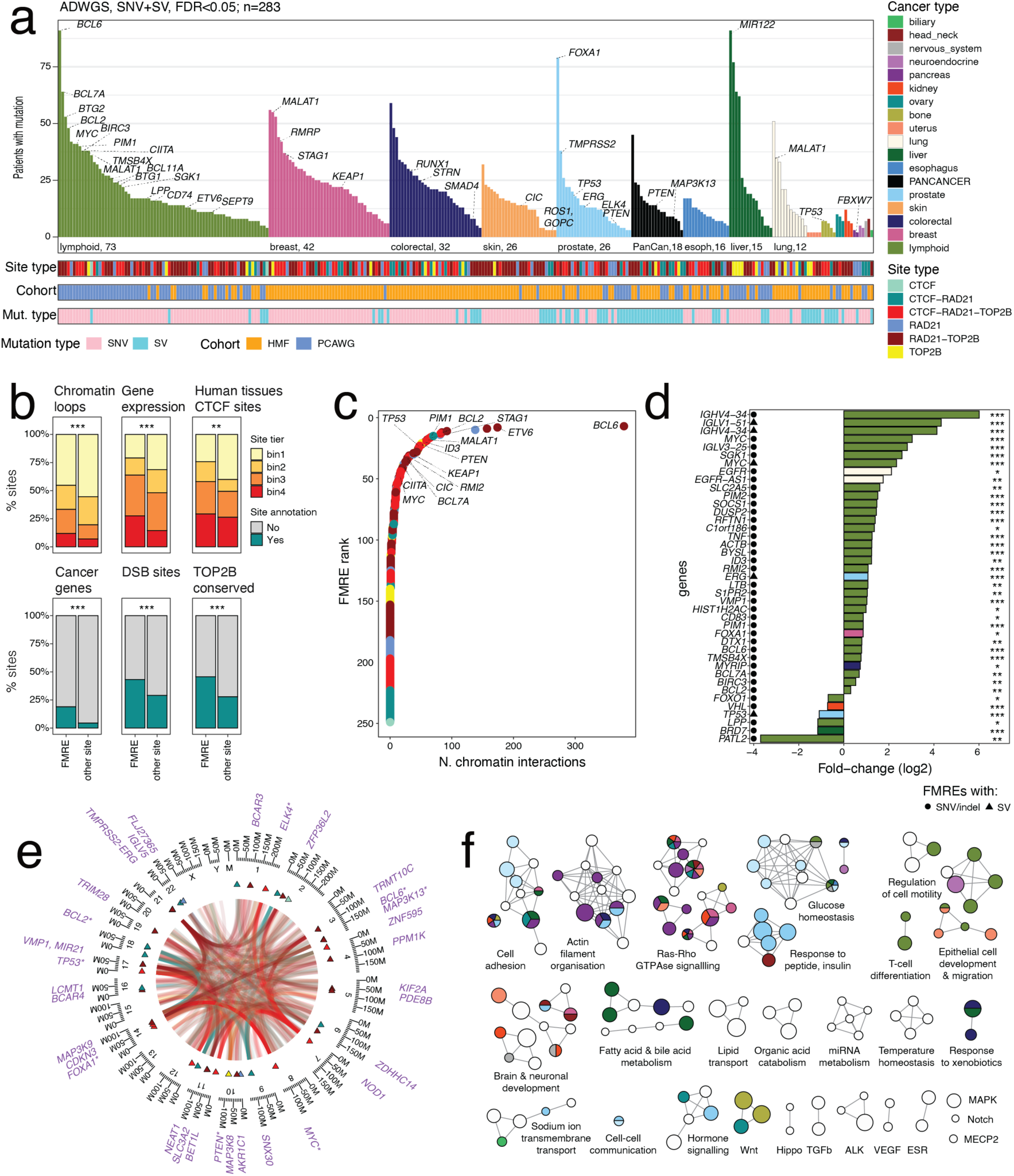
Frequently mutated regulatory elements (FMREs) at TOP2B binding sites. **(a-b)** FMREs are binding sites of TOP2B, CTCF, or RAD21 that have enriched SNVs/indels or SVBPs compared to a null model derived from flanking genomic sequences (ActiveDriverWGS, FDR < 0.05). FMREs were predicted separately for cancer types and primary and metastatic samples. Known cancer genes with non-coding mutations are highlighted. **(b)** Functional characteristics of FRMEs. Compared to other TCRBS sites, FMREs are enriched in highly transcribed elements with higher evolutionary conservation and chromatin interactions. P-values from hypergeometric tests or Wilcoxon tests are shown. **(c)** FMREs ranked by the number of chromatin interactions from 27 human tissue types. Top genes with FMREs are labelled. **(d)** Mutations in a subset of FMREs associate with differential expression of target genes. Symbols indicate associations identified from SNVs and indels (circles) or SVBPs (triangles). P-values are from ANOVA analyses and account for gene CNA for SNV/indel based FMREs (FDR < 0.05). **(e)** Circos plot of translocations at FMREs combined from primary cancers and metastases. Known cancer genes at FMREs are shown. **(f)**. Enrichment map of biological processes and molecular pathways enriched in non-coding FMREs integrated across primary and metastatic cancers (ActivePathways, FDR < 0.05). The network shows enriched pathways as nodes in which similar pathways that share many genes are connected by edges. Colors indicate the cancer types in which the pathways were identified.

We mapped the epigenomic characteristics of FMREs. Most FMREs were bound by TOP2B- CTCF-RAD21 or TOP2B-RAD21, more than expected from our catalogue of binding sites (hypergeometric P = 9.1 x 10^-8^). Functional annotations were also enriched, such as conserved TOP2B binding in mouse liver (P = 2.6 x 10^-9^), *in vitro* evidence of DSBs (P = 1.9 x 10^-6^), and constitutive CTCF binding in human cell types (P = 0.0014) (**Figure 4b**). Thus, TOP2B binding at FMREs explains the mutational enrichments beyond CTCF or RAD21 binding alone. FMREs were associated with higher gene expression (P = 6.7 x 10^-9^, Wilcoxon test) and promoter-enhancer interaction frequency (P = 3.4 x 10^-5^), indicating their putative roles in gene regulation and chromatin architecture in cancer. Some FMREs appeared as hotspots of three-dimensional chromatin interactions: 142/253 (56%) FMREs had at least one promoter-enhancer interaction and 32 FMREs (13%) at least 25 interactions across 27 human tissue types. These chromatin interaction hubs often occurred at cancer driver genes such as *PTEN*, *TP53*, *BCL6* and *ETV6* (**Figure 4c**). Considering distal FMREs though promoter-enhancer chromatin interactions highlighted 21 additional distally-located cancer driver genes including *CHEK2* and *CCND3* that were not apparent when annotating FMREs and genes linearly (**Figure S4a-c**). Thus, some TOP2B-associated FMREs may regulate cancer genes via promoter-enhancer interactions and topologically associating domains ^25^.

We evaluated the functional importance of FMREs using complementary evidence from cancer transcriptomes and pathway information. First, we asked if mutations or SV breakpoints in FMREs were associated with differential expression of target genes in matching cancer transcriptomes. We found 41 genes with significant differential expression (FDR < 0.05 from ANOVA and F-tests) (**Figure 4d**), including FMRE mutations at *FOXA1* in breast cancer, *VHL* in kidney cancer, *TP53* in prostate cancer, as well as several associations with lymphoma driver genes. Next, we asked whether FMREs with enriched non-coding mutations showed pathway-level evidence of cancer relevance. To that end, we distinguished a subset of FMREs that were not dominated by protein-coding mutations (203/244) and performed an integrative pathway enrichment analysis ^55^ that prioritised FMREs and genes found in multiple cancer types. We found 176 significantly enriched GO biological processes and Reactome ^56^ pathways that were indicative of hallmark cancer processes such as cell adhesion, cell motility and migration, epithelial cell development and signalling pathways of MAPK, VEGF, Notch Hippo, and Ras- Rho GTPases, among others (ActivePathways FWER < 0.05, **Figure 4f**). These data suggest that non-coding FMREs converge on biological pathways across cancer types.

### TOP2B binding sites at established mutational hotspots

We examined FMREs at known cancer genes, most of which were detected due to significant enrichments of SNVs and indels in multiple cancer types (FDR < 0.05, ActiveDriverWGS) (**Figure 5a**). First we focused on mutational hotspots and loss-of-function mutations that often affect oncogenes and tumor suppressors, respectively ^57^, and found 15 genes with five or more such mutations in FMREs (**Figure 5b**). First, *IDH1* mutations in the R132H/C/L hotspot were associated with TOP2B-RAD21 binding and were enriched in biliary and central nervous system cancers (**Figure S5a**). Mutations in isocitrate dehydrogenase (IDH) genes define a lower-risk subtype of glioma and drive a hypermethylator phenotype ^58,59^. Secondly, the *VHL* gene encoding the von Hippel-Lindau tumor suppressor protein was highlighted due to a TOP2B- RAD21 bound FMRE that was enriched in loss-of-function mutations of frameshift indels and nonsense SNVs in 61 kidney cancers in PCAWG and HMF (2.7 tumors expected, FDR < 10^-17^ from ActiveDriverWGS) (**Figure S5b**). *VHL* mutations were associated with a significant down- regulation of the gene in matching cancer transcriptomes, potentially through nonsense-mediated decay (FDR = 1.8 x 10^-4^, F-test) (**Figure 4d, Figure S5e**). Thirdly, the T790M mutational hotspot in epidermal growth factor receptor (EGFR) occurred at a TOP2B-RAD21-bound FMRE that was identified in metastatic lung cancer (42 tumors mutated *vs*. 10 expected; FDR = 1.3 x 10^-9^) (**Figure S5c**) and associated with *EGFR* upregulation (FDR = 0.026) (**Figure 4d, Figure S5e**). EGFR T790M mutations cause acquired resistance to tyrosine kinase inhibitors ^60^. Some known cancer driver genes were also enriched in non-coding mutations at TOP2B binding sites. The pioneer factor *FOXA1* ^61^ included a triple site in the final exon and 3’UTR that was enriched in coding and non-coding mutations in breast and prostate cancers (ActiveDriverWGS, FDR ≤ 10^-12^) (**Figure S5d**). The FMRE was enriched in indels (114/220 indels *vs*. 22 expected; binomial P = 3.0 x 10^-55^) that caused rearrangements of the 3’UTR of *FOXA1* and frameshift mutations in the last exon. These examples suggest that the DNA breakage and repair activity of TOP2B may contribute to positively selected driver mutations in cancer.

**Figure 5.**
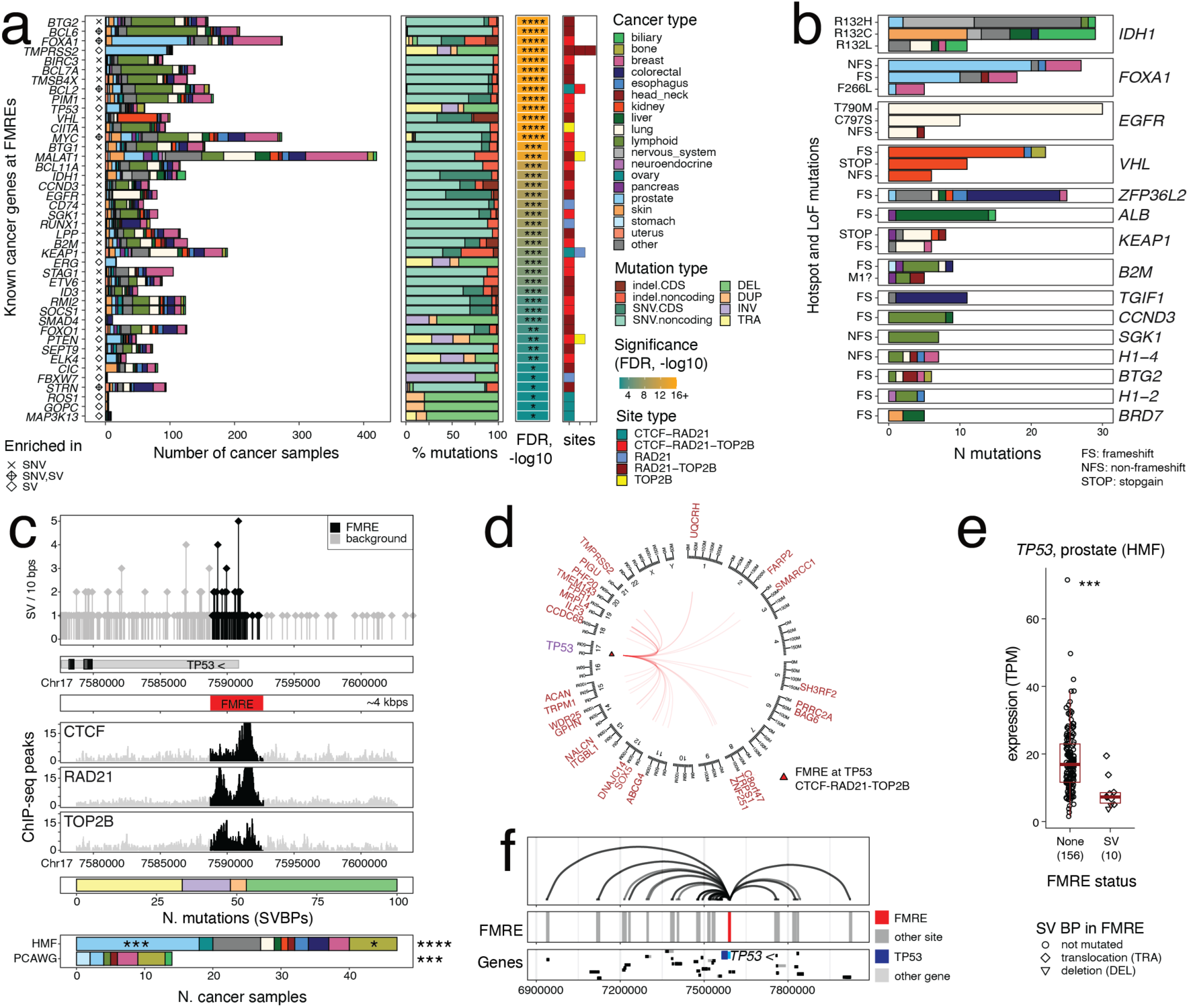
TOP2B binding sites at established cancer driver mutations. **(a)** List of known cancer driver genes at FMREs identified due to enriched SNVs and indels or SV breakpoints in the driver analysis (Fig. 4). The stacked barplots show numbers of primary and metastatic cancer samples affected at each FMRE, types of mutations at each FMRE, and types of TOP2B binding sites. **(b)** TOP2B binding occurs at oncogenic mutational hotspots and loss-of-function mutations in known driver genes. Mutations (≥ 5) in the affected genes are grouped by individual amino acid substitutions or impact as frameshift (FS), non-frameshift (NFS), or nonsense mutations (*i.e.*, STOP). Colors indicate cancer types and mutation counts are shown across primary and metastatic cancers. (**c-f**) TOP2B-CTCF-RAD21-bound FMRE at the promoter of the tumor suppressor gene *TP53* is enriched in SV breakpoints (SVBPs). **(c)** Genomic locus of the FMRE at the promoter of *TP53*. Shown are the SVBP frequencies in the FMRE (∼4 kbps; black) and the adjacent flanking sequence in grey (top), DNA-binding signals of CTCF, RAD21, and TOP2B as ChIP-seq peaks in black (middle), and the counts of SVBPs and the affected cancer samples, colored by alteration types and cancer types (bottom). The FMRE was independently identified in two pan-cancer cohorts as well as metastatic prostate cancer and bone cancer (asterisks show FDR-values from ActiveDriverWGS). **(d)** Circos plot visualisation of translocations at the FMRE located at the *TP53* promoter. The FMRE at chr17 is indicated with a triangle and arcs extending to other chromosomes show translocations with the FMRE. Adjacent putative target genes are labelled (brown). **(e)** In metastatic prostate cancer, SVBPs in the FMRE associate with reduced expression of *TP53* in matched cancer samples, indicating that SVs in this TOP2B-bound element disrupt tumor suppressor activity of TP53. FDR-adjusted P-values from ANOVA analysis are shown. **(f)** The FMRE at the *TP53* promoter has many long-range chromatin interactions. Arcs show chromatin interactions identified from 27 types of normal human tissues.

### FMREs with structural variation hotspots

We reviewed the FMREs with SV hotspots at known cancer genes. One of the most prominent FMREs occurred in the promoter of the tumor suppressor gene *TP53*. This triple site was enriched in pan-cancer SV breakpoints in both PCAWG and HMF cohorts that affected 61 cancer samples (6.3 expected, min FDR = 4.1 x 10^-25^ from ActiveDriverWGS) (**Figure 5c**). The FMRE was also found in metastatic prostate cancer. It predominantly involved deletions and translocations with various loci genome-wide (**Figure 5d**). Analysis of matching transcriptomes revealed that the subset of metastatic prostate cancers with SVBPs at the FMRE showed significant reduction in *TP53* expression compared to cancers lacking the SVBPs (FDR = 4.4 x 10^-5^) (**Figure 5e**). This suggests a TOP2B-mediated recurrent rearrangement of the *TP53* promoter that leads to transcriptional inhibition of tumor suppression in cancer. The FMRE is also a hotspot of promoter-enhancer chromatin interactions (**Figure 5f**), suggesting a mechanistic explanation where TOP2B-mediated resolution of genomic entanglements from 3D chromatin interactions could lead to DNA damage and positively selected structural variants at the tumor suppressor.

Several other recurrent SV hotspots were found at TOP2B binding sites. First, the *TMPRSS2*- *ERG* locus involved three adjacent TOP2B-RAD21 FMREs in *TMPRSS2* and another intronic triple site in *ERG* that were highly significant in the driver analysis and collectively affected 105 prostate cancer samples in the two cohorts (8.9 expected, min FDR = 6.6 x 10^-36^) (**Figure S6a-c**). *TMPRSS2-ERG* gene fusions are hallmark drivers of prostate cancer ^62^ and it has been shown previously that their formation involves TOP2B-induced DNA DSBs ^22^, lending confidence to our findings. As expected, these SVs associated with significant increase in *ERG* expression in matching prostate cancer transcriptomes (*FDR* = 1.8 x 10^-4^) (**Figure S6d**). Secondly, the *MYC* oncogene included a frequently altered triple site in the promoter and 5’ UTR with SVs in 14 B-cell non-Hodgkin’s lymphomas in PCAWG (0.3 expected, FDR = 3.2 x 10^-14^) that encoded translocations with immunoglobulin heavy chain genes at chr14, a known driver mechanism in lymphoma ^63^ (**Figure S7a-b**). Accordingly, lymphomas with these SVs showed substantial *MYC* upregulation compared to samples lacking these rearrangements (FDR = 8.9 x 10^-11^) (**Figure S7c**). The FMRE also had multiple promoter-enhancer interactions (**Figure S7d**), exemplifying the roles of TOP2B binding in genome architecture and structural variation in cancer. Thirdly, the most significant FMRE with SVs was found at the *NOD1* gene in metastatic colorectal, esophageal and lung cancers with SVBPs in 49 cancer samples (1.2 expected; FDR = 1.2 x 10^-54^), including many translocations genome-wide (**Figure S8**). *NOD1* encodes an intracellular sensor of microbial components involved in innate immunity ^64^ and the recurrent SVs highlight a putative driver role triggered by structural variation. These data suggest that TOP2B-driven DSBs may lead to structural rearrangements of cancer driver genes.

### Non-coding mutations at the *RMRP*/*CCDC107* FMRE drive cancer *in vivo*

We asked if TOP2B binding sites include non-coding cancer driver mutations. Analysis of TOP2B binding sites in this large dataset of whole cancer genomes revealed dozens of tissue- specific FMREs with enriched somatic mutations in breast, prostate, colorectal cancers as well as lymphomas and melanomas (**Figure 4a-b**). The functional and epigenomic annotation, evolutionary conservation and pathway associations of these FMREs suggest the presence of previously uncharacterised cancer driver mutations affecting gene regulation and chromatin architecture.

We examined one top mutational hotspot located on chr9 at the non-coding RNA *RMRP* that we identified as a strong triple site co-bound by TOP2B, CTCF and RAD21 in our ChIP-seq data (**Figure 6a**). It was identified as a putative non-coding driver element in 42 metastatic breast cancers (ActiveDriverWGS, FDR = 1.4 x 10^-8^). While only detected in the breast cancer dataset, this element was also highly mutated in other cancer types and showed at least 5% mutation frequency in breast, stomach, skin, ovary, and kidney cancers, with 197 samples in total. Thus, the FMRE at the *RMRP* locus is a non-coding mutation hotspot with a mutation frequency above 2% in two pan-cancer cohorts. Interestingly, most mutations in the FMRE (210/215) were non- coding (**Figure 6b-c**). This mutational hotspot has been described in prior cancer genomics studies with *in vitro* evidence of functionality ^66^. However, recent pan-cancer reports suggest that these are potentially passenger mutations generated through local mutational processes ^24^. In addition to the non-coding RNA gene *RMRP* with mitochondrial function, the FMRE occurs at the 5’ end of *CCDC107* that encodes a little-studied membrane protein, and the 3’ UTR of *ARHGEF39* that encodes a Rho guanine nucleotide exchange factor.

**Figure 6.**
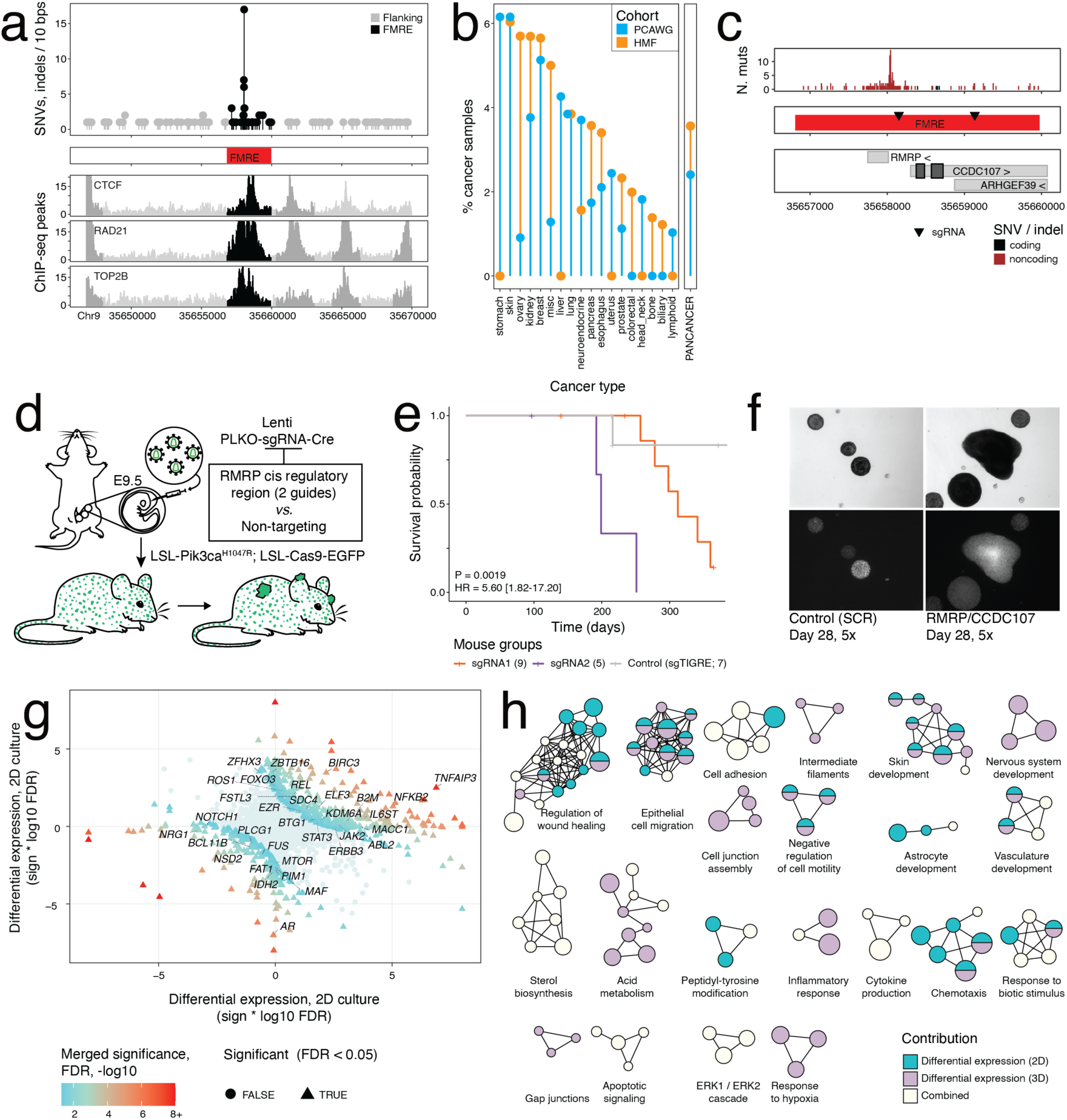
Experimental validation of the FMRE at the RMRP/CCDC107 locus. **(a)** FMRE at a triple site at the *RMRP* and *CCDC107* locus is enriched in SNVs and indels in breast cancer. SNV and indel counts from the primary cancers (PCAWG) and metastases (HMF) are shown. DNA-binding patterns of TOP2B, CTCF, and RAD21 from ChIP-seq experiments are shown at the bottom. (b) The FMRE at the *RMRP/CCDC107* locus is frequently mutated in multiple cancer types. Mutation frequencies of SNVs and indels are shown for primary cancers and metastases. The average pan-cancer mutation frequency is shown on the right. **(c)** Higher-resolution view of the *RMRP/CCDC107* FMRE in the human genome with the mutational hotspot (top), the FMRE and sgRNAs shown as triangles (middle), and the genes *RMRP*, *CCDC107*, and *ARHGEF39* at the site. Most mutations in the FMRE are non-coding (brown). **(d)** Overview of experimental validation. Two sgRNAs targeting the orthologous mouse FMRE were delivered by lentivirus using an ultrasound-guided *in utero* injection into mouse embryos. In parallel, two sgRNAs of the human FMRE were delivered into MCF10A human mammary epithelial cell lines. **(e)** Genome editing of the *RMRP* FMRE locus causes earlier tumor onset *in vivo*. Kaplan-Meier curve shows tumor-free survival of the mice with targeted and control sgRNAs. P-values and hazard ratios (HR) were derived using Cox proportional-hazards regression and Wald test. HRs with 95% confidence intervals are shown. (f) Genome editing of the *RMRP* FMRE locus leads to enhanced growth of MCF10A cells. Three-dimensional growth assay (Matrigel) compares control (SCR)-treated cells (left) and cells with mutations in the *RMRP/CCDC107* FMRE (right). Cells were imaged on the 28-day timepoint. **(g)** Differential gene expression analysis of FMRE-mutant MCF10A cells from transcriptome-wide comparisons with SCR-treated cells. Integrative analysis of the RNA-seq data prioritizes genes that are jointly either up-regulated or down-regulated in 2D and 3D growth assays of FMRE-targeted cells. The scatterplot shows differential expression scores from 2D assays (X-axis) and 3D assays (Y-axis), with positive and negative scores showing FDR-weighted up- regulation and down-regulation in FMRE-mutant cells, respectively. Significantly deregulated genes are colored (merged FDR< 0.05 from DPM ^65^) and known cancer genes from the Cancer Gene Census database are labelled. **(h)** Genes deregulated in FMRE-mutant cells are enriched in invasion-related and developmental processes. The enrichment map is a network-based visualisation that shows significantly enriched pathways from the integrative analysis of 2D and 3D growth assays (ActivePathways, FWER < 0.05). Enriched pathways are shown as nodes in the network in which similar pathways that share many genes are connected by edges. Nodes are colored based on the statistical evidence from either 2D or 3D growth assays or both, or are painted white if the pathway was only identified by integrating 2D and 3D experiments. Node size corresponds to the number of genes per pathway.

We sought to validate the role of this FMRE experimentally using functional genomics approaches in a human mammary epithelial cell line and a skin cancer mouse model. To that end, we performed CRISPR/Cas9 genome editing of the locus in MCF10A human breast epithelial cells using one sgRNA targeting the major mutational hotspot in the *RMRP* promoter and another sgRNA targeting the non-coding regions of *CCDC107* and *ARHGEF39*. In parallel, we edited the conserved orthologous FMRE in the mouse genome using ultrasound-guided *in utero* injection of two independent sgRNAs into the surface ectoderm of mouse embryos, expressing the sensitizing Pik3ca^H1047R^ oncogene ^67^ (**Figure 6d**). In this skin cancer mouse model, FMRE mutations triggered significantly earlier tumor onset compared to scramble control sgRNAs (P = 0.0019, Cox proportional-hazards test) (**Figure 6e**). In MCF10A cells, FMRE mutations caused enhanced proliferation phenotypes in a three-dimensional (3D) growth assay (**Figure 6f**). To complement these functional assays, we performed transcriptome-wide sequencing of FMRE- edited MCF10A cells from 2D and 3D growth conditions and asked which genes were differentially expressed relative to scramble-treated control cells (**Methods**). Directional integration of transcriptomics data using the DPM method ^65^ revealed 641 genes that were either up-regulated or down-regulated jointly in the two growth conditions and by both sgRNAs (FDR < 0.05) (**Figure 6g**). Pathway enrichment analysis of differential gene expression patterns confirmed the phenotypes of FMRE-mutant cells we observed: the major themes involved cell migration and motility, regulation of wound healing, cell adhesion, and several developmental processes (**Figure 6h**). Interestingly, the genes *RMRP*, *CCDC107* and *ARHGEF39* adjacent to the FMRE were not differentially expressed in FMRE-mutant cells; however, several gene-regulatory and signalling genes with cancer functions such as *NOTCH1*, *FAT1*, *ERBB3*, *TNFAIP3* and *NFKB2* were differentially expressed (**Figure S9**). Known cancer genes were overall more frequent than expected from chance alone (hypergeometric P = 0.0051). Non-coding driver mutations at this FMRE may have complex epigenetic and regulatory effects that can be deciphered by further examining the downstream deregulation of the transcriptome.

Collectively, these experiments characterise the frequently mutated *RMRP* locus as a non-coding cancer driver and exemplify the binding patterns of TOP2B in cancer genomes as a rich resource for follow-up experiments.

## DISCUSSION

While TOP2B-mediated DSBs are essential in regulating chromatin architecture during transcription, development, and differentiation, these DSBs can also be a double-edge sword, potentially leading to genomic rearrangements and mutations in cancer. In this study, we undertook a comprehensive approach by integrating a genome-wide map of TOP2B binding in human cancer samples with a vast dataset comprising thousands of whole cancer genomes, spanning primary tumors to metastatic lesions. Our findings reveal that TOP2B binding sites are subject to the influence of local mutational processes, which give rise to SNVs, indels and structural rearrangements, both in primary cancers and metastatic chemotherapy-treated cancers. The notable enrichment of SVBPs at TOP2B binding sites is consistent with our understanding of the role of TOP2B in generating DSBs and aligns with previous reports linking TOP2B to chromosomal translocations ^12–14,21,22^, which lends confidence to our TOP2B binding map.

However, the frequent occurrence of SNV and indel mutations at TOP2B binding sites is somewhat unexpected within the context of topoisomerases and may be indicative of the complex processes involved in DSB formation and repair. Recently, rare protein-coding mutations in TOP2A have been linked to a mutational indel signature in cancer genomes, demonstrating the capability of topoisomerases to generate indel mutations ^68^. Besides somatic mutations in cancer genomes, these processes may exert a broader influence on the evolutionary dynamics of the germline genome.

The most striking patterns of somatic mutations emerge precisely at triple and double binding locations where TOP2B co-binds DNA with CTCF and/or RAD21. It is noteworthy that a pronounced enrichment of mutations at CTCF and cohesin binding sites has been observed in multiple cancer types ^31,69–71^. However, our study goes a step further by demonstrating that TOP2B binding at these sites offers a plausible explanation for these mutational patterns compared to the co-factors alone, suggesting that TOP2B-mediated DNA cleavage might be a key source of mutations. Moreover, our findings expose intriguing associations with gene expression and chromatin interactions that point to context- and tissue-specific determinants of TOP2B-mediated mutagenesis in primary tumors, as well as uncover hotspot regions of chemotherapy induced mutagenesis, warranting future investigation. The evolutionary conservation and widespread activity of these sites underscore their functional significance and lend strong support to the notion that TOP2B-mediated mutagenic processes may play a pivotal role in cancer initiation, progression, and heterogeneity as well as therapy resistance.

The prominent presence of TOP2B across the cancer genome encompasses both well-established driver mutations, as well as hotspot regions harboring frequent genomic rearrangement at dozens of cancer driver genes in primary and metastatic tumors. Remarkably, TOP2B-associated SVBPs converge on hotspots associated to oncogene activation, such as the FMREs observed at *MYC* and *TMPRSS2-ERG*, or conversely, abolishing tumor suppressive pathways, as exemplified by the FMRE at the *TP53* promoter. TOP2B’s influence extends to recurrent mutation hotspots in driver genes like *VHL*, *FOXA1*, *EGFR*, and *CDH1*, where the protein-coding SNVs and indels cause aberrant loss or gain of function. Intriguingly, our TOP2B binding map also highlights hundreds of FMREs harboring putative non-coding driver mutations. Some of these sites exhibit additional evidence of altered regulation of target genes, frequent long-range chromatin interactions, or changes in DNA motifs. FMREs often display evolutionary conservation, overlap with transcriptionally active genes and promoter-enhancer interactions, and many are likely to hold functional roles in cancer. As a proof of concept, the direct causal role of TOP2B in the generation of structural variants at the *TMPRSS2* and *ERG* loci and production of *de novo* driver fusions of prostate cancer has been established by a prior extensive mechanistic study ^22^, implicating TOP2B as the driver of these FMREs. We report TOP2B binding at a diverse set of FMREs suggesting that TOP2B may contribute to the genesis of driver mutations and their subsequent positive selection in cancer more commonly than documented to date. However, unraveling the intricacies of positive selection in cancer and fully understanding the mutagenic processes at TOP2B binding sites will require substantial further research.

We generated *in vitro* and *in vivo* validation data on one FMRE, a prominent non-coding mutational hotspot at the non-coding RNA gene *RMRP*. Our *in vivo* data of tumor initiation and *in vitro* data of enhanced cellular growth in the three-dimensional growth condition highlight this locus as a bona fide non-coding cancer driver. This element is highly mutated in multiple cancer types and cohorts, lending confidence to its broad pan-cancer roles. However, it appears as a region with complex gene-regulatory interactions that do not control adjacent gene expression but instead regulate numerous genes transcriptome-wide, many of which are involved in cancer hallmark processes and key signal transduction pathways. Thus, more work is needed to decipher the mechanisms of non-coding mutations and molecular cancer phenotypes.

Our study has several limitations. First, our profiling of TOP2B binding was conducted on a relatively limited number of clinical samples from hepatocellular carcinoma, which provides only a partial view of TOP2B activity in cancer. Assessing TOP2B binding in human cells has been experimentally challenging to date. We profiled this tissue type by extending our previous robust experimental protocols in mouse liver ^3^. While our binding maps represent liver cancer, most cancer types analysed consistently revealed mutational enrichments at TOP2B binding sites. These further exhibited evolutionary conservation and were constitutively active, suggesting a broader, pan-tissue activity of TOP2B. Secondly, apart from triple sites, we observed mutational enrichments at CTCF-RAD21 sites, which also displayed evolutionary conservation for CTCF or TOP2B binding. We speculate that these sites include TOP2B binding sites in other cell or tissue types, which may not have been fully captured in our profiles due to the inherent cancer heterogeneity. Thirdly, our mutational analysis entails two large and diverse cancer genomics datasets of untreated primary cancers and treated metastatic cancers, respectively, that have variable sample sizes and differences in data processing. Notwithstanding, it is worth noting that our findings are consistently replicated across multiple cancer types and cohorts. Despite these challenges, our TOP2B binding maps, along with annotations of mutational processes and known and putative driver mutations, represent a valuable resource for computational and experimental follow-up studies.

The mutational processes occurring at TOP2B binding sites may be initiated by an array of factors, such as TOP2B poisoning by chemotherapeutic drugs, dietary components, or environmental compounds. Subsequent processing of protein-trapped DSB ends can contribute to chromosomal translocations and the development of secondary malignancies, as documented in leukemia ^10,14^. Furthermore, potentially tumorigenic DNA lesions and translocations can originate from TOP2B-DNA transactions during endogenous cellular processes such as RNA Pol II pause release and transcriptional elongation, as shown in prostate and breast cancers ^21,22^. Our study underscores the prevalence of small somatic mutations and chromosomal rearrangements in both primary cancers and metastatic chemotherapy treated cancers at TOP2B binding sites, which appear to be more common than previously recognized. Future mechanistic studies and integrative analyses bridging molecular cancer profiles with clinical and lifestyle data will be essential for advancing our understanding of the role of TOP2B in cancer mutagenesis and etiology, and for uncovering the risks and off-target effects of widely used TOP2-targeting chemotherapies.

In summary, we posit that TOP2B functions as a double-edged sword in cancer biology. While it plays a critical role in relieving topological stress and maintaining genome integrity, it also paradoxically serves as a source of somatic cancer mutations. Our comprehensive catalogue of FMREs at TOP2B binding sites, harboring both known and putative driver mutations, stands as a valuable resource for the formulation of novel hypotheses and mechanistic studies.

## METHODS

### Tissue collection

Three liver hepatocellular carcinoma (HCC) samples were collected from patients who underwent liver transplantation at the Ajmera Transplant Centre of Toronto General Hospital (ON, Canada) in 2016-2019. The HCC samples were from male patients with mean age 59 (age range 56-64) and of hepatitis C etiology. All three samples were of stage II (T2N0M0) and moderately differentiated histologic grade. Written informed consents were obtained from all participants prior to the procedures according to the guidelines of Multi-Organ Transplant Biobank (Toronto, ON, Canada). The study was approved by the Research Ethics Boards of University Health Network and University of Toronto, Toronto, ON, Canada.

### Chromatin immunoprecipitation (ChIP-seq) experiments

Approximately 30 mg of HCC tumor sample was used for each ChIP reaction. Frozen HCC tissues were dounce homogenized into single cell suspensions and cross-linked with 1% formaldehyde in Solution A (50 mM Hepes–KOH, 100 mM NaCl, 1 mM EDTA, 0.5 mM EGTA) for 20 min at RT. Fixation was quenched by adding glycine to a final concentration of 125 mM. After two washes with ice cold PBS, cells were filtered through 100 μm cell strainer to remove any connective tissue. Cell lysis and nuclei isolation were carried out in 10 mL of LB1 buffer (Diagenode) by rotating 20 min at 4 C, followed by nuclear lysis in 5 mL of LB2 (Diagenode) and rotating 10 min at 4C. For chromatin shearing, the cell pellet was resuspended in Shearing Buffer (Diagenode) supplemented with complete proteinase inhibitors (Roche), and sonicated for 5 cycles (30 s ON, 30 s OFF) with the Bioruptor Pico sonicator (Diagenode). Chromatin was cleared by centrifugation (13,000 rpm for 10 min at 4 C). For ChIP, chromatin lysates were combined with 5 ug of anti-TOP2B, anti-CTCF, or anti-RAD21 antibodies and incubated overnight in IP buffer (Diagenode) rotating at 4 C, and then with 70 μl of pre-blocked (0.5 mg/mL BSA) Dynabeads protein G (ThermoFisher) for 4 hours. Beads were then washed six times with RIPA buffer (50 mM Hepes–KOH, pH 7.5; 500 Mm LiCl; 1 mM EDTA; 1% NP-40 or Igepal CA-630; 0.7% Na–Deoxycholate), and one time with TBS (20 mM Tris–HCl, pH 7.6; 150 mM NaCl), and resuspended in ChIP Elution buffer (50 mM Tris–HCl, pH 8; 10 mM EDTA; 1% SDS). Cross- linking was reversed by overnight incubation at 65 C. Cellular proteins and RNA were digested with Proteinase K (Invitrogen) and RNaseA (Ambion). ChIP DNA was purified with P:C:I method. ChIP-seq libraries were prepared using NEBNext Ultra II DNA Library preparation Kit following the manufacturer’s instructions (NEB, E7645). Libraries were amplified for 9 cycles using Q5 Hot Start High-Fidelity DNA Polymerase (NEB, M0493L), purified and size-selected with AMPure XP PCR purification beads (Beckman Coulter, A63881), and quantified with 2100 Bioanalyzer (Agilent). Input control DNA extracted from sonicated cell lysates of each sample were processed in parallel. Libraries were sequenced with Illumina NovaSeq SP in a paired-end configuration. Unprocessed ChIP-seq data are available in ArrayExpress (E-MTAB-13301).

### ChIP-seq data analysis and defining TCRBS

We generated four libraries of binding sites for CTCF, three sets for RAD21, and two sets for TOP2B. Quality assessment of sequencing data was performed using the FastQC method ^72^. Sequences were aligned to the human genome hg19 using the Bowtie2 method (v2.1.0) with the parameters “-1” and “-2” for paired-end reads ^73^. The Samtools method (v1.2) was used to sort, clean, and format the data ^74^. Peak calling was performed in individual replicates separately using the MACS2 method ^75^ with standard parameters except for the parameter “-f BAMPE” to specify paired-end reads and the parameter “-p 1e-3” to initially establish a lenient p-value threshold. As output, narrowPeak files including the standard BED columns with four additional columns were generated, including signal values, P-values, q-values, and relative summit positions. Multiple testing correction was performed using the Benjamini-Hochberg (BH) False Discovery Rate (FDR) method ^76^. TCRBS were first filtered for significance and fold-change (FDR < 0.05, FC ≥ 2). TCRBS in chromosomes 1-22, X, and Y were selected and TCRBS in unmappable regions of the genome were excluded, as defined in the ENCODE project for hg19 (ENCFF001TDO) ^41^. Based on the distribution of TCRBS widths, TCRBS lengths were initially fixed to 550 bps around summits. Within each factor (*e.g.*, all CTCF libraries), we then merged the overlapping coordinates and calculated the median summit position and q-value. For CTCF and RAD21, we selected only the TCRBS where binding was detected in at least two different libraries. Due to technical limitations, we used a more sensitive process for TOP2B binding sites and selected sites if binding was detected in at least one of the two libraries. As tumors are heterogenous and TOP2B ChIP-seq is an experimentally sensitive procedure with specific challenges in human cells, we opted for this approach to extract a wider set of TOP2B binding sites. TOP2B binding sites were filtered stringently similarly to RAD21 and CTCF sites (FDR < 0.05; FC ≥ 2). To define sites bound by the three factors TOP2B, CTCF, or RAD21, we extended each site using a lenient width of ±1 kbps around each site midpoint and merged the resulting sites based on sequence overlaps. Seven types of sites were defined based on the binding of the three factors: TOP2B-CTCF-RAD21 (*i.e.*, triple sites), TOP2B-RAD21 (*i.e.*, double sites), CTCF-RAD21 sites, TOP2B-only sites, RAD21- only, sites CTCF-only sites, and CTCF-TOP2B sites. Relatively few CTCF-only and CTCF- TOP2B sites were found and these were excluded from most downstream analyses.

### Somatic SNVs, indels, and SVBPs in whole cancer genomes

Somatic SNVs and indels in whole cancer genomes were retrieved from two datasets, PCAWG ^77^ and HMF ^47^, that were in hg19. Hypermutated samples with >90,000 somatic SNVs or indels were excluded. For PCAWG, the consensus dataset of SNVs and indels was used. For HMF, variant calls were further filtered as follows. We excluded data from multiple tumors from the same patient by selecting the first tumor ID alphabetically and removed samples lacking sufficient informed consent. Variant calls in HMF VCF files were filtered (filter = pass). Cancer types in PCAWG and HMF were consolidated to 18 cancer types based on organs and anatomical sites.

Consolidated cancer types with at least 25 samples in both HMF and PCAWG cohorts were analysed separately, while the pan-cancer analyses also included all other cancer samples and types in the two cohorts. We mapped the functional effects of SNVs and indels on protein-coding genes using the ANNOVAR software ^78^ and RefSeq gene definitions. Mutational signatures of single base substitutions (SBS) were compiled for the two cohorts separately. SBS signatures of the PCAWG dataset were retrieved from the consensus dataset by Alexandrov *et al*. (2020) ^37^.

HMF SBS signatures were computed using the SigProfiler software with SigProfilerMatrixGenerator (v1.2) and SigProfilerExtractor (v1.1.4) ^79^ using SBS signatures of the COSMIC database (V3) ^79,80^. Each SNV was assigned to the most probable SBS signature given its trinucleotide sequence. All indel mutations were combined for signature analyses.

Somatic structural variants (SVs) from the PCAWG and HMF datasets were analysed separately. For PCAWG, we used the consensus SV dataset developed by Li *et al*. (2020) ^38^. For HMF, SVs identified using the gridss software ^81^ were further filtered using recommended steps to select high-quality SVBP calls (filter = PASS, qual ≥ 1000, AS > 0, RAS > 0) and high-confidence SV events for which both SVBPs were available were selected. Common SV annotations were assigned: duplications (DUP), deletions (DEL), inversions (INV), and translocations (TRA). We also mapped SVBPs to genes using ±3 kbps flanking regions. For differential gene expression analysis described below, we also computed gene copy-number alterations (CNAs) via somatic CNA segments in PCAWG and HMF, assigning the median total copy number to each gene based on overlapping CNA segments.

### Cancer transcriptomes

For the PCAWG dataset, gene expression data was retrieved from the consensus transcriptomics dataset with fragments per kilobase per million values (FPKM-UQ)^46^. For the HMF dataset, gene expression data was processed from paired fastq files, with reads aligned using trimGalore and Cutadapt (version 0.6.6) ^82^, expression quantified as transcripts per million (TPM) values using the Kallisto method (v0.46.2) ^83^, and gene-level TPM counts imported using tximport and tximportData (v1.18.0). Low-confidence transcriptomes with less than 35% pseudoalignment were excluded. We also excluded samples for which WGS data was not available. For patients with multiple cancer transcriptomes, we selected the samples with the highest level of tumor purity.

### Long-range chromatin interactions (loops)

We annotated TCRBS using long-range chromatin interactions (*i.e.*, chromatin loops) from promoter-capture HiC experiments in 27 human tissues from the study by Jung et al. (2019) ^48^ in which higher-confidence interactions were selected (freq > 10; -log10_score > 3). To annotate TCRBS to genes through chromatin loops, we assigned one loop anchor to a TCRBS using sequence overlap, and the other loop anchor to a transcription start site (TSS) of a protein-coding gene. TSSs were retrieved from Ensembl BioMart for GRCh37. From these distal interactions of sites and genes we excluded those interactions that were already identified based on site proximity to gene (i.e., within ±3 kbps). We focused on long-range chromatin interactions involving FMREs and visualised these as networks in Cytoscape.

### Genomic and epigenomic annotations of TCRBS

TCRBS were annotated using genomic, epigenomic, and functional features. First, we analysed the overlaps of TCRBS with 162,000 CTCF binding sites in 70 human cell lines in the ENCODE project ^41^. TCRBS were classified by CTCF binding as tissue-specific (binding in <10% of cell lines), intermediate, or constitutively active (binding in >90% of cell lines). Evolutionary conservation of CTCF binding sites in five mammalian livers and conservation of TOP2B binding in mouse liver were retrieved from the study by Uusküla-Reimand *et al.* (2016) ^3^. Chromatin states of TCRBS were obtained from the core 15-state model of the Epigenomics Roadmap project ^45^ in ten normal human tissues (breast, colorectum, esophagus, kidney, liver, lung, ovary, pancreas, skin, stomach) that represented the most frequent cancer types in the WGS datasets we analysed (see below). Experimentally mapped DSB sites from two cell lines (Mcf7, Nalm6) were retrieved from the study by Canela *et al*. (2017) ^4^ and merged based on overlapping coordinates. Experimentally mapped common fragile sites (CFS) from three cell lines (HeLa, HS68, U2OS) were retrieved from the study by Macheret *et al*. (2020) ^49^, mapped to hg19 from hg38 using the LiftOver software ^84^, and merged based on coordinate overlaps. Next, we used the GENCODE database to annotate TCRBS to protein-coding genes, lincRNA genes, antisense, miRNA, and sense-intronic non-coding genes, as well as immunoglobulin (IG) and T-cell receptor (TR) genes. These genes were also used to filter the transcriptomics datasets of PCAWG and HMF using Ensembl gene identifiers. TCRBS were associated with adjacent genes based on ±3 kbps flanking windows. The TCRBS with no adjacent genes were considered intergenic. CTCF binding motifs in TCRBS were identified using the FIMO method ^85^ of MEME software (version 5.4.1) using the HOCOMOCOv11_core_HUMAN TF motif dataset and default parameter settings.

### Grouping TCRBS by gene expression and chromatin looping activity

We grouped TCRBS into four bins on their functional activity separately for gene expression and long-range chromatin interactions, as follows. Binning was performed for each class of TCRBS separately (triple, double, CTCF-RAD21, RAD21-only, TOP2B-only sites). Sites lacking chromatin loops or intergenic sites with no adjacent gene expression within ±3 kbps were assigned to bin 1 (*i.e.*, the no-activity bin). The three remaining bins of equal size represented sites with low, medium, and high activity based on gene expression levels, or chromatin loop counts, respectively. For chromatin interactions, we counted how many loops each TCRBS overlapped across all tissues in the dataset, such that the highest counts were assigned to the TCRBS with many interactions in many different tissues. For gene expression, we selected the most highly expressed gene per site for bin assignment. Bins were assigned for each cancer type and cohort using the median expression values. This analysis was limited to the cancer types for which at least five matching transcriptomes were available. In addition, TCRBS were grouped to bins by median pan-cancer expression values in HMF and PCAWG.

### Statistical analysis of TCRBS annotations

Evolutionary conservation and constitutive activity of TCRBS was assessed using one-tailed hypergeometric tests to check if sites with high evolutionary conservation (five species) or constitutive CTCF binding (>90% of human tissues) were enriched in a type of TCRBS relative to other TCRBS combined. Similarly, we asked if DSBs, CTCF motifs, and conserved TOP2B binding sites in mouse liver were enriched. Next, we evaluated the enrichments of chromatin states by comparing TOP2B-bound TCRBS with control TCRBS lacking TOP2B binding (triple *vs*. CTCF-RAD21 sites, double *vs*. RAD21-alone sites). Using two-tailed hypergeometric tests, we evaluated the 15 core chromatin states in ten types of human tissues and excluded states with few sites (<250) and reported statistically significant results as odds-ratio values (FDR <0.05). We then studied TOP2B binding in the context of pan-cancer gene expression in PCAWG and HMF as well as individual cancer types in the two cohorts. Median gene expression values were log1p-transformed and converted to Z-scores.

TCRBS were assigned to the adjacent gene (±3 kbps) with the highest gene expression value (Z-score). The Z-scores were compared using non-parametric tests (Wilcoxon rank-sum tests) between TCRBS bound and not bound by TOP2B, and classes of TCRBS with and without TOB2B binding (*i.e.*, triple *vs*. CTCF-RAD21 sites, double *vs*. RAD21-alone sites). The proportions of intergenic TCRBS (*i.e.*, at >3 kbps of genes) with respect to TOP2B binding were analysed using hypergeometric tests. Third, we studied TOP2B binding in the context of promoter-enhancer chromatin loops pooled across a panel of normal human tissues. For each TCRBS, we counted the number of overlapping loop anchors and used non-parametric analysis with Wilcoxon rank-sum tests to compare TCRBS with and without TOP2B binding, as described above.

### Mutational processes of SNVs and indels in TCRBS

We studied the mutational processes of SNVs and indels at TOP2B binding sites using the RM2 method, which compares the mutation burden in a set of genomic elements with their flanking regions given trinucleotide and megabase-level covariation using negative binomial regression ^32^. We used the default RM2 window size of 100 bps such that sites of 200 bps around the midpoints of TCRBS (i.e., ±100 bps) were compared with controls of 200 bps flanking sequences up- and downstream of TCRBS. For an unbiased comparison of site sets of different sizes, RM2 was run in down- sampling mode that repeatedly analysed a fixed number of sites from the overall site pool for 100 iterations and selected the results corresponding to the median P-value. First, we analysed different types of TCRBS (triple, double, CTCF-RAD21, TOP2B-alone, RAD21-alone), excluding CTCF-only and CTCF-TOP2B sites with few instances. RM2 down-sampled to 10,000 sites to approximate the least frequent site type. The 18 cancer types and the pan-cancer sets for PCAWG and HMF were analysed separately. First, we analysed the classes of TCRBS in all cancer types, adjusted all results for multiple testing, and filtered (FDR < 0.05). Grassy hills plots in RM2 were used for visualisation. Second, we studied the four bins of TCRBS grouped by gene expression or chromatin looping activity. Gene expression analysis was conducted for cancer types and cohorts for which at least five samples of the same cancer type with transcriptomics data were available. RM2 down-sampled the analysis to 1000 TCRBS to approximate the smallest bins of TCRBS. While reducing bias from unbalanced site sets, this more stringent down-sampling rendered the analysis relatively underpowered compared to others. A combined BH FDR correction was applied across the expression- and chromatin looping-based analyses, significant findings were selected (FDR < 0.05) and were visualised as a heatmap by ordering the results from expression and chromatin looping analyses by directional associations. Third, RM2 was used to compare TCRBS with three classes of functional annotations: double-strand break (DSB) sites, common fragile sites (CFS), and sites with conserved TOP2B binding in mouse livers relative the same types of TCRBS lacking these genomic annotations. RM2 down-sampled to 1,000 TCRBS to reduce biases from sample sizes. Significant results were selected after multiple testing correction (FDR < 0.05). Lastly, we asked if SNVs of specific mutational signatures of single base substitutions (SBS) were enriched in TOP2B binding sites. SBS mutation frequencies in sites with and without TOP2B binding (TOP2B-CTCF-RAD21 vs. CTCF-RAD21 sites; TOP2B-RAD21 vs RAD21 sites) were compared using one-sided hypergeometric tests and significant results were reported (FDR < 0.05).

### Mutational processes of structural variants in TCRBS

We analysed SVBPs in TCRBS using two complementary methods. First, we counted the numbers of SVBPs per site in the different types of TCRBS. The few sites bound by CTCF-only and CTCF-TOP2B were excluded. As additional genome-wide control sites, we used the set of non-overlapping 2-kbps genomic windows that excluded all TCRBS Poorly-defined gap regions from the UCSC database (UCSC_hg19_gap.txt.gz) were also excluded from controls. We compared per-site SVBP counts in different types of TCRBS using Wilcoxon rank-sum tests by counting pan-cancer SVs in PCAWG samples, HMF samples, and combined samples of the two cohorts. To associate SVBP burden with gene expression, we also compared four bins of TCRBS grouped by pan-cancer gene expression. This analysis only included TOP2B-bound TCRBS (*i.e.*, triple, double, and TOP2B-alone sites). SVBP counts in sites in the highest bin of sites (Bin 4, *i.e.*, most gene expression) were compared to the three lower bins (bins 1-3 combined) using Wilcoxon rank-sum tests. Sites grouped by chromatin looping activity were analysed similarly. To group TCRBS by gene expression, sites were assigned to the adjacent gene (± 3kbps) with the highest median value of pan-cancer expression. For pan-cohort analyses, the cohort with the higher value was used. The pan-cancer expression analyses were also replicated in cancer type specific analyses by comparing the SV counts and the transcriptomes of cancer samples of matching cancer types. Lastly, we also asked if SVBP burden in TOP2B-bound TCRBS was associated with functional genomic annotations including DSBs, common fragile sites (CFS), and evolutionary conservation of TOP2B or CTCF binding in mouse livers. This analysis only included TOP2B-bound TCFBS and compared per-site SVBP counts in TCRBS that had or lacked specific functional annotations (*i.e.*, CFS, DSB, conservation) using Wilcoxon rank-sum tests.

### Analysis of SVBPs in TCRBS relative to SNV/indel burden

Having established that SNVs and indels were enriched in TCRBS, we analysed SVBP burden relative to SNV/indels in TCRBS. SVBPs were compared in TOP2B-bound TCRBS relative to non-TOP2B bound TCRBS in two comparisons: triple sites *vs*. CTCF-RAD21 sites, and double sites *vs*. RAD21- only sites. One-tailed hypergeometric tests determined whether SVBPs in TOP2B-bound sites were more frequent than expected from non-TOP2B bound sites given the distributions of SVBPs and SNVs/indels in TCRBS. The analysis was repeated for all cancer types in PCAWG and HMF. All P-values were adjusted for multiple testing and significant results were selected (FDR < 0.05). Results with at least 10 SVBPs were visualised, and FDR values were capped (FDR < 10^-16^).

### Discovery of frequently mutated elements (FMREs) as cancer drivers

We performed a driver discovery analysis to find individual TCRBS that had frequent somatic mutations (*i.e.*, FMREs) using the ActiveDriverWGS method ^25^. ActiveDriverWGS utilises Poisson regression to find genomic regions with increased mutation burden relative to adjacent genomic background sequence and its trinucleotide-level sequence content. Simple somatic mutations (SNVs, indels) and SVBPs were analysed separately. Within these two groups, samples of the cancer types and each of the two cohorts (PCAWG, HMF) were also analysed separately. Binding sites with zero mutations were assigned a conservative P-value of 1.0. A stringent multiple testing correction using BH FDR was conducted across cancer types and the two cancer cohorts and significant results were selected (FDR < 0.05). SNVs/indels and SVBPs were analysed separately since SVBPs were less frequent genome-wide and different parameter settings of ActiveDriverWGS were used. For SNVs/indels, ActiveDriverWGS used the default background sequence around TCRBS (±50 kbps). For SVBPs, two adaptations were implemented: (i) a wider, more conservative background sequence around TCRBS was used to account for the lower genome-wide SVBP frequency for a more accurate background mutation rate (± 500 kbps), and (ii) additional pan-cancer analyses of SVBPs in PCAWG and HMF cohorts were performed for increased statistical power. Multiple testing correction and selection of SVBP-based FMREs was similar to the SNV/indel analysis. FMREs with translocations were visualised as circos plots using the ggbio package in R ^86^. Known cancer genes associated with FMREs were retrieved from the COSMIC Cancer Gene Census database ^87^ (downloaded July 31^st^ 2021). FMREs were also visualised in the context of promoter-enhancer chromatin interactions pooled from a panel of normal human tissues ^48^. We compared FMREs and non-FMRE TCRBSs in the context of chromatin loop counts, adjacent gene expression levels, and number of cell lines with CTCF binding using Wilcoxon tests, and their location at known cancer genes, DSB sites, and conserved TOP2B binding sites using hypergeometric tests.

### Associating mutations in FMREs with target gene expression

To evaluate the functional roles of mutations in FMREs, we asked if these associated with gene expression within ±3 kbps. Each cancer type was analysed separately, as were PCAWG and HMF samples, and SNVs/indels and SVBPs. First, we selected genes for which at least five mutations in FMREs and matching RNA-seq data were available and did not show low overall expression (mean expression ≥ 1).

Gene expression values were log1p-transformed. For the SNV/indel analysis, we excluded samples with high gene amplifications (CN > 8) for more conservative estimates of expression associations. To analyse differential expression, we compared gene expression values of FMRE- mutated and non-mutated tumor samples using linear regression with the gene CN as a covariate, and derived P-values from F-tests using ANOVA. For increased stringency, multiple testing correction with BH FDR was conducted across all tests combining the cancer types in HMF and PCAWG. Significant results were selected (FDR < 0.05). For SV BPs in FMREs, the gene expression analysis was performed similarly, except that gene CN values were not used to filter samples or in covariates, since SVBPs included CNAs. To analyse *ERG* gene expression in the context of TMPRSS2/ERG structural variants, we combined SVBPs from the TMPRSS2 FMRE and the *ERG* FMRE and compared *ERG* expression in metastatic prostate cancers with and without these SVBPs.

### Pathway enrichment analysis of non-coding FMREs

An additional analysis of FMREs with non-coding SNVs/indels was performed for the pathway analysis and to confirm that the regions were not only dominated by protein-coding mutations. To this end, we re-defined TCRBS by subtracting protein-coding (CDS) exons and analysed these again with ActiveDriverWGS. The pathway enrichment analysis of FMREs with frequent non-coding SNVs/indels integrated mutational enrichments across all cancer types we analysed. To map pathways to TCRBS, we first mapped TCRBS to protein-coding genes, selecting the most significant TCRBS to every protein-coding gene using a ±3 kbps flanking window around genes. To integrate signals across cancer types, the most significant FMRE per gene was selected for each cancer type from either PCAWG or HMF, resulting in a matrix of top P-values of protein-coding genes for the 18 cancer types. We used integrative pathway enrichment analysis of the ActivePathways method ^55^ by merging evidence across the cancer types, prioritising genes that showed mutational enrichments in TCRBS in multiple cancer types. ActivePathways also included sub-significant TCRBS and therefore considered mutational enrichments beyond the list of FMREs. All protein-coding genes were used as the background set, gene sets of 25-500 genes were included in the analysis, significant pathway enrichments were selected using the Holm family-wise error rate (FWER < 0.05) and default parameters were used otherwise. Pathway analysis considered functional gene sets representing molecular pathways of the Reactome database ^88^ and GO biological processes that were retrieved from the g:Profiler ^89^ web site (2023-10-06). The results were visualised as an enrichment map using the Cytoscape software (v3.10.0). We curated the map manually to identify major functional themes ^90^.

### Lentivirus production and transduction for *RMRP* FMRE experiments

Large-scale production and concentration of lentivirus were performed as previously described ^67,91^. Briefly, 293T cells (Invitrogen R700-07) were seeded on a poly-L-lysine coated 15 cm plates and transfected using PEI (polyethyleneimine) method in a non-serum media with lentiviral construct of interest (pLKO-Cre sgRNA v4; Addgene plasmid 158032) along with lentiviral packaging plasmids psPAX2 and pPMD2.G (Addgene plasmids 12259, 12260). 8 hours post-transfection media was added to the plates supplemented with 10% Fetal bovine serum and 1% Pencillin-Streptomycin antibiotic solution (w/v). 48 hours later, the viral supernatant was collected and filtered through a Stericup-HV PVDF 0.45-μm filter, and then concentrated ∼2,000-fold by ultracentrifugation in an MLS-50 rotor (Beckman Coulter).

### CRISPR guides for *RMRP* FMRE experiments

For MCF10A cell culture experiments we used two guides: ATAGGCTTTCAGAGGCATTG and CTAGAGTTCCAGATATGAAG. For mouse *in utero* transductions in the skin we used two guides: AAGTATCATGCCTAAAACAA and CCCGGCTACCTGTAAAATGA. The NTC sequence is GAAGGAGGCTACACCCGTTA.

### In utero lentiviral transduction for *RMRP* FMRE experiments

Ultrasound-guided lentiviral injection and related procedures have been described ^67,91^. Briefly, to deliver the lentiviral sgRNAs targeting gene of interest, a non-invasive, ultrasound-guided in utero injection method was employed, which selectively transduces single-layered surface ectoderm of living E9.5 mouse embryos in a clonal fashion.

### 2D cell culture for RNA extraction for *RMRP* FMRE experiments

75,000 sgRNA-infected C3 MCF10A cells ^92^ were plated into four 12-well plates (Sarstedt 83.3921). The cells were grown for 24 hours and scraped into 100ul of RNA Lysis buffer (Zymo Research R1051), then stored in -80C prior to RNA extraction.

### 3D sphere formation assay for *RMRP* FMRE experiments

For three-dimensional sphere formation experiments, MCF10A cells were plated on Matrigel Matrix (Corning 354234) as described previously ^93^. The initial starting count of 15,000 cells were plated on 24-Well Plates (Sarstedt 83.3922) precoated with 100ul of Matrigel, and grown in MCF10a Assay Media, made with DMEM/F12 (Wisent 319-084-CL), 2% Horse Serum (Gibco 16050122), 0.5ug/ml Hydrocortisone (Sigma-Aldrich H0888), 100ng/ml Cholera Toxin (Sigma-Aldrich C8052), 10ug/ml Insulin (Sigma-Aldrich I5500), 1x Pen/Strep (Wisent 450-201-EL), 5ng/ml EGF (Wisent 511-110-EU) and 2% Matrigel. Media was changed every other day for three weeks.

### Sphere dissociation from Matrigel for RNA extraction for *RMRP* FMRE experiments

Media was removed and plates were placed on ice for 5 minutes. 1ml of cold PBS was added to each well and contents were transferred into microtubes on ice. A second wash was performed with 800ul and the remaining contents were added into their respective microtubes. Samples were spun at 5 min, 4C at 1000 x g. The supernatant was removed, and the cell pellets were washed two more times with cold PBS. After the final spin, each cell pellet was resuspended in 200ul Zymo Lysis buffer (Zymo Research R1051).

### Total RNA sequencing for *RMRP* FMRE experiments

RNA samples were extracted from cells using the Zymo Quick-RNA Microprep Kit (Zymo Research R1051). Three replicates were used for each of two targeted sgRNAs and the scramble control sgRNA. After extraction, sample integrity was checked with RNA Qubit (Invitrogen Q32852) and the Bioanalyzer. Total RNA sequencing with single-end reads at 40 million reads per sample was performed at the core facility of the Lunenfeld-Tanenbaum Research Institute.

### Analysis of RNA-seq data from *RMRP* FMRE experiments

We uniformly processed RNA-Seq single-end reads for 18 samples using the Rsubread ^94^ method (v2.12.0). Reads were aligned to the human genome using default parameters using the reference genome hg38 from the Ensembl database (downloaded 2024-01-16). Read counts for each gene defined in the Gencode gtf annotation file (v45, downloaded 2024-04-23) were obtained. We filtered genes with low expression values (<10 counts in 70% of samples). Differential gene expression analysis was performed on filtered genes with raw counts using the edgeR ^95^ method (v4.0.16). Gene expression was compared separately for 2D and 3D cultures by combining two sgRNA treatments of three replicates each and the scramble-treated control of three replicates.

Differential gene expression was tested using a quasi-likelihood negative binomial generalized log-linear model (glmQLFit) implemented in edgeR. P-values were adjusted for multiple testing using the FDR method and significant genes were selected for volcano plots (FDR < 0.05).

Directional pathway enrichment analysis was designed to jointly consider transcriptomic changes across both invasion assays and prioritise genes and pathways that differentially expressed in both experiments in the same direction (*<u>i.e.</u>*, sign of fold-change). Pathway analysis was performed with the directional ActivePathways method ^55^ (v2.0.3). Genes with low expression were removed from the background gene set for the pathway analysis. Gene Ontology and Reactome gene sets were downloaded from the g:Profiler ^89^ website (downloaded 2024-06-14).

Pathway enrichment analysis considered gene sets of 25 to 500 genes and significant results were selected using the Holm multiple-testing correction (FWER < 0.05). The results were visualized as an enrichment map using the Cytoscape software (v3.9.1). We curated the map manually to identify major functional themes ^90^.

## Acknowledgments

This work was supported by the Canadian Institutes of Health Research (CIHR) Project Grant PJT-162410 to J.R., and the Investigator Award to J.R. from the Ontario Institute for Cancer Research (OICR), and the New Investigator Award of the Terry Fox Research Institute (TFRI) to J.R. M.D.W. was supported by the Canada Research Chairs Program. L.U.R. was supported by a Next Generation of Scientists Award from the Cancer Research Society (PIN25558), the Estonian Research Council fellowship, NSERC grant (RGPIN-2019-07014) to M.D.W., as well as a Cancer Research Society/Isaiah 40:31 Memorial Fund Operating grant to M.D.W. and J.R. Funding to OICR is provided by the Government of Ontario. The results shown here are in whole or part based upon data generated by the TCGA Research Network: https://www.cancer.gov/tcga. We acknowledge the contributions of the many clinical networks of ICGC and TCGA who provided samples and data to PCAWG. We thank the patients and their families for their participation in ICGC and TCGA projects. This publication and the underlying study have been made possible partly based on the data that Hartwig Medical Foundation has made available to the study.

## Author contributions

C.A.L. and J.R. analyzed the data. L.U.R. and S.A.A. performed the ChIP-seq experiments. L.U.R., C.A.L., and J.R. interpreted the data and wrote the manuscript. L.U.R. and M.D.W. designed and supervised the ChIP-seq experiments. R.O., E.L., R.S., and D.S. performed the functional validation experiments. N.A., Z.P.K., K.C.L.C., D.A.R., and H.H. contributed to data analysis. E.P. and M.B. contributed clinical samples. J.R., M.D.W., and L.U.R. conceived the project. J.R. supervised the project. The authors reviewed and edited the manuscript and approved the final version.

