## Supplementary Figures for "Topoisomerase IIb binding underlies frequently mutated elements in cancer genomes"

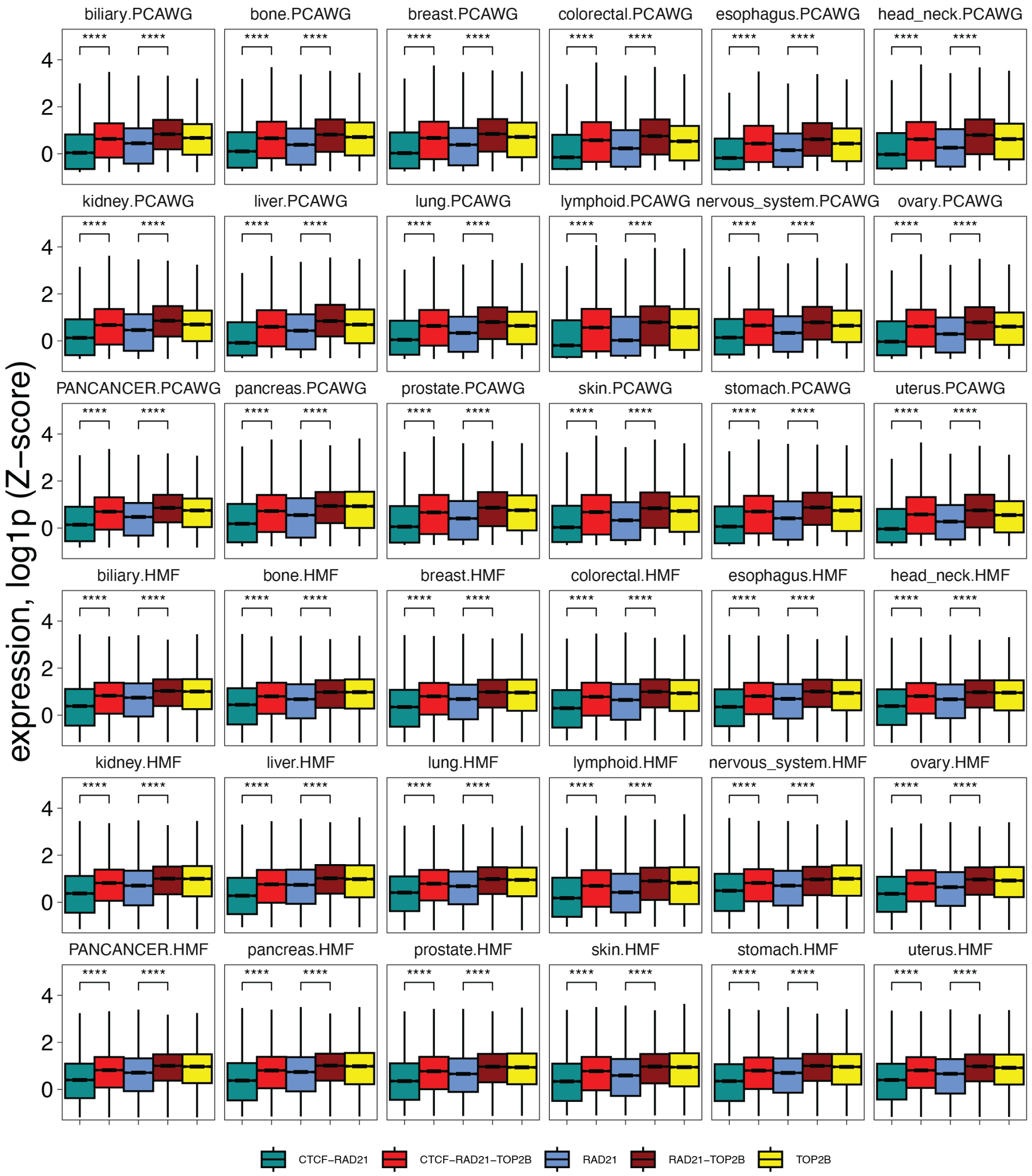


**Figure S1. Associations of TOP2B binding and gene expression. (a)** TOP2B binding sites associate with higher gene expression in multiple cancer types. Matching site types with and without TOP2B binding were compared (triple sites *vs*. CTCF-RAD21 sites, double sites *vs*. RAD21 sites) using nonparametric analysis with Wilcoxon rank-sum tests (asterisks). RNA-seq data from PCAWG and HMF were used for corresponding cancer types. For each site, the adjacent gene (±3 kbps) with the highest median expression was used with Z-scores of log-transformed transcription quantities (FPKM-UQ for PCAWG or TPM for HMF). Neuroendocrine tumors were excluded due to limited RNA-seq data. (*, $P$ <0.05; **, $P < 0.01$; ***, $P < 0.001$; ****, $P < 2.2 \times 10^{-16}$).

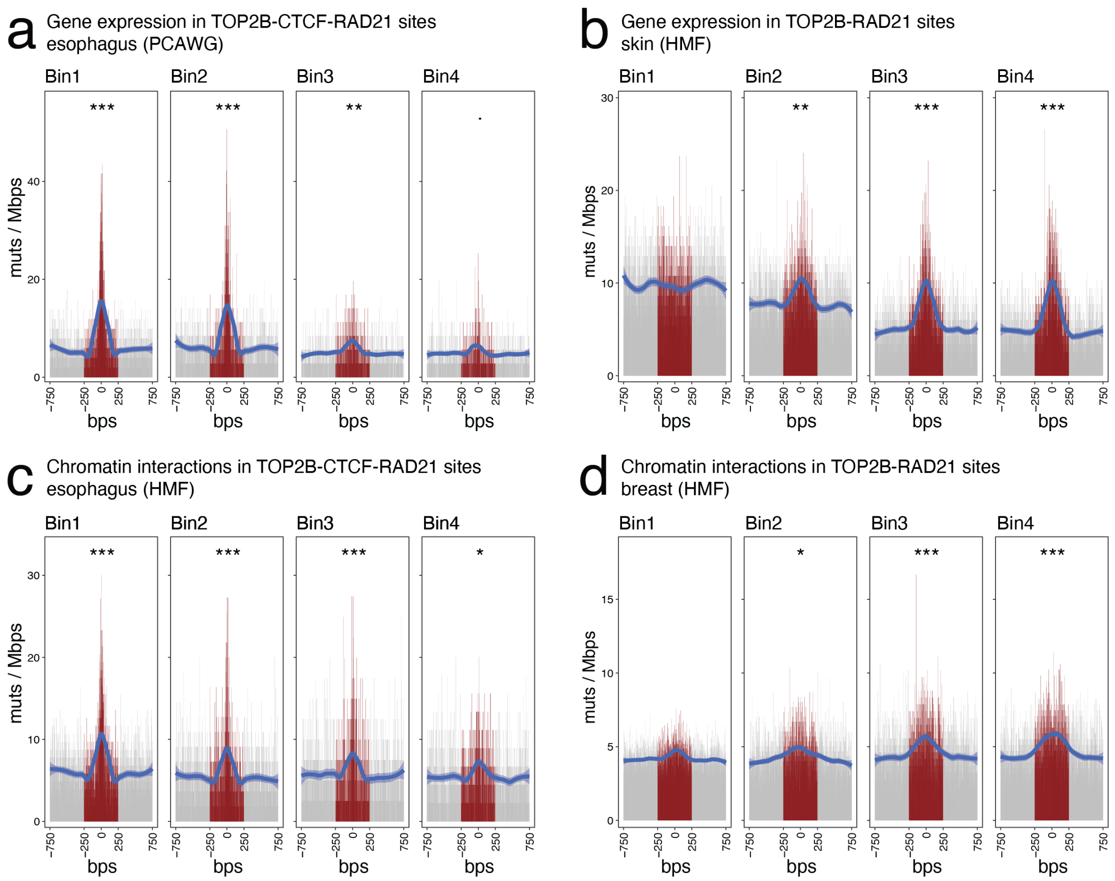


**Figure S2. Context-specific local mutational processes at TOP2B binding sites.** RM2 analysis of local mutational processes in triple sites (TOP2B-CTCF-RAD21) and double sites (TOP2B-RAD21) grouped by functional activity, *i.e.,* mRNA expression of adjacent genes (panels (a, b)) and promoter-enhancer chromatin interactions (i.e., chromatin loops) (right; panels (c, d)). Binding sites were grouped into four bins (left to right) representing sites with no activity (bin1), and sites representing low, medium and high activity (bins 2,3,4) represent equal bins having 1/3 of sites. For gene expression analysis, we used transcriptomes of relevant cancer types from the PCAWG and HMF datasets. For chromatin looping analysis, promoter-enhancer interactions from multiple normal human tissues and cell lines were retrieved from the study by Jung *et al.* (2019). **(a)** Higher gene expression associated with lower local mutation burden at triple sites in esophageal cancer. **(b)** Higher gene expression associated with higher local mutation burden at double sites in skin cancer. **(c)** Higher chromatin looping activity associated with lower local mutagenesis in triple sites in esophageal cancer. **(d)** Higher chromatin looping activity associated with higher local mutagenesis in double sites in breast cancer.

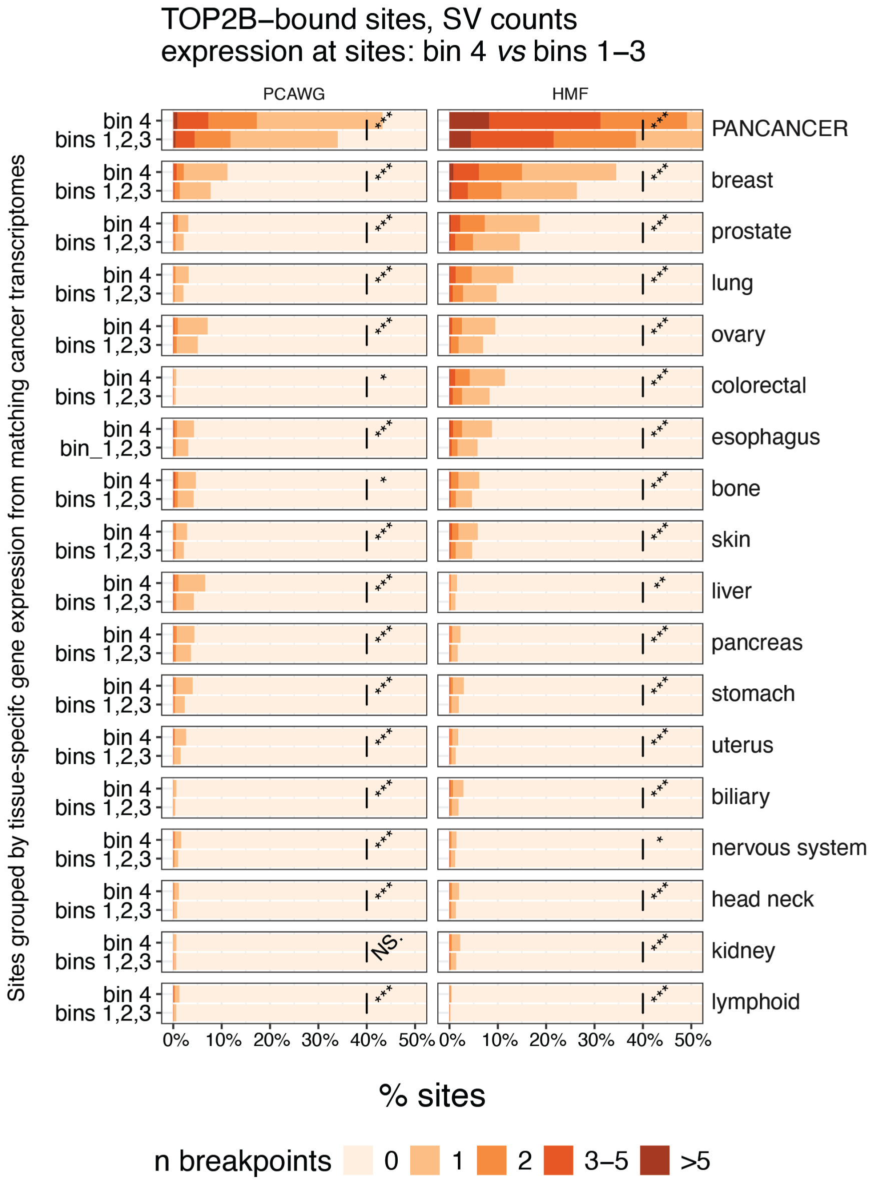


**Figure S3. Mutagenesis of SV breakpoints (SVBPs) in TOP2B binding sites associates with higher gene expression.** We compared TOP2B-bound sites at highly expressed genes (bin 4) with TOP2B-bound sites at genes with low or medium expression or intergenic sites (bins 1-3). The analysis included all sites bound by TOP2B (triple, double, TOP2B-only) SVBPs counts per site group were compared using Wilcoxon rank-sum tests (*, $P$ <0.05; **, $P < 0.01$; ***, $P < 0.001$; ****, $P < 2.2 \times 10^{-16}$). Stacked bar plots show the fractions of TOP2B-binding sites grouped by SVBPs in the sites.

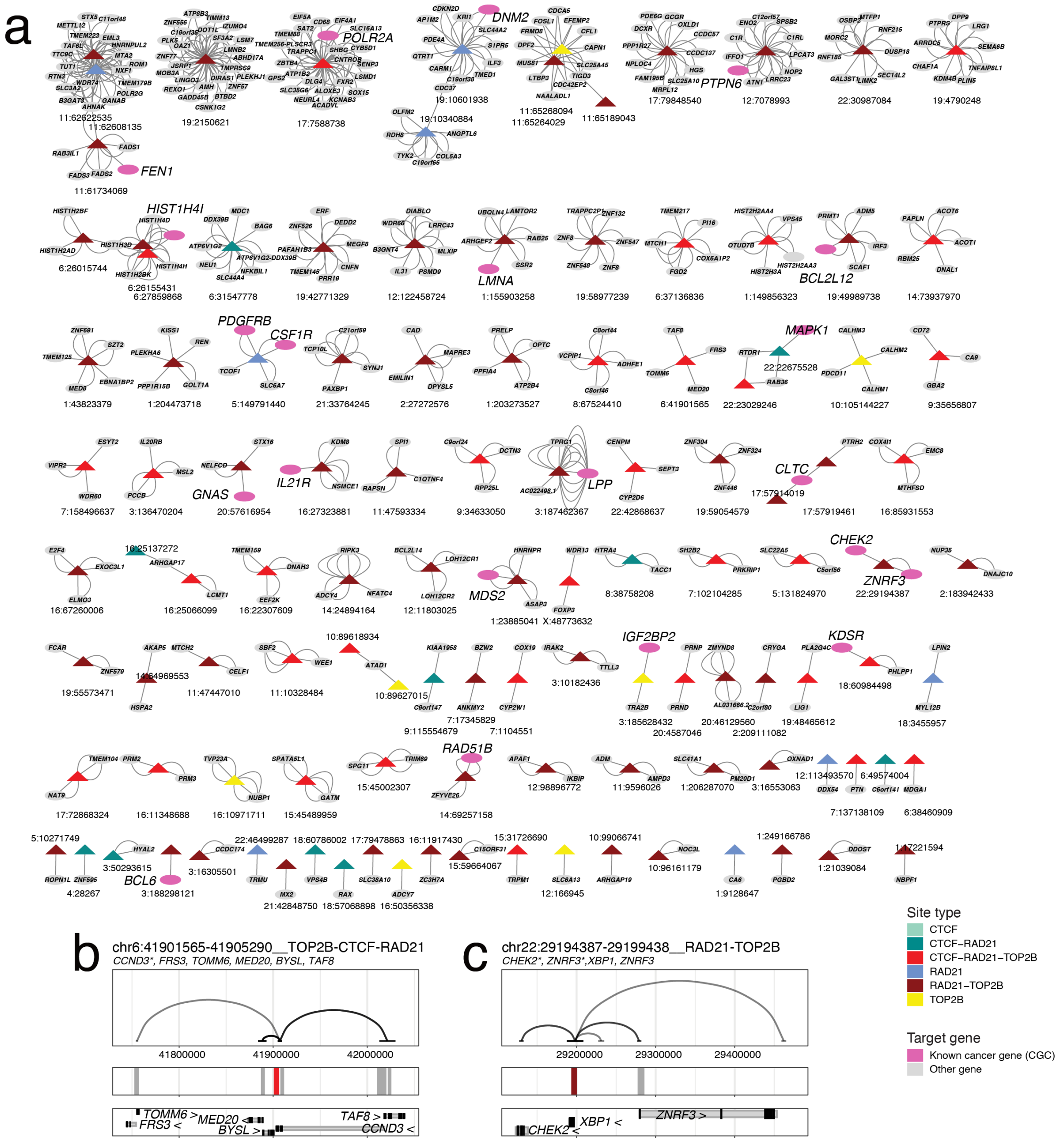


**Figure S4. Promoter-enhancer chromatin interactions at FMREs. (a)** All FMREs that have promoter-enhancer interactions (chromatin loops) to gene transcription start sites (TSSs) as identified in promoter-capture HiC datasets from Jung *et al.* (2019). FMREs are shown as triangles, circles represent genes and edges represent chromatin loops. Known cancer genes are shown in hot pink. **(b-c)** Examples of FMREs with frequent promoter-enhancer chromatin interactions at known cancer genes (*CCND3*, *CHEK2*). Facets show relevant genomic annotations: promoter-enhancer chromatin interactions (top); FMREs and non-significant sites (middle); and adjacent protein-coding genes.

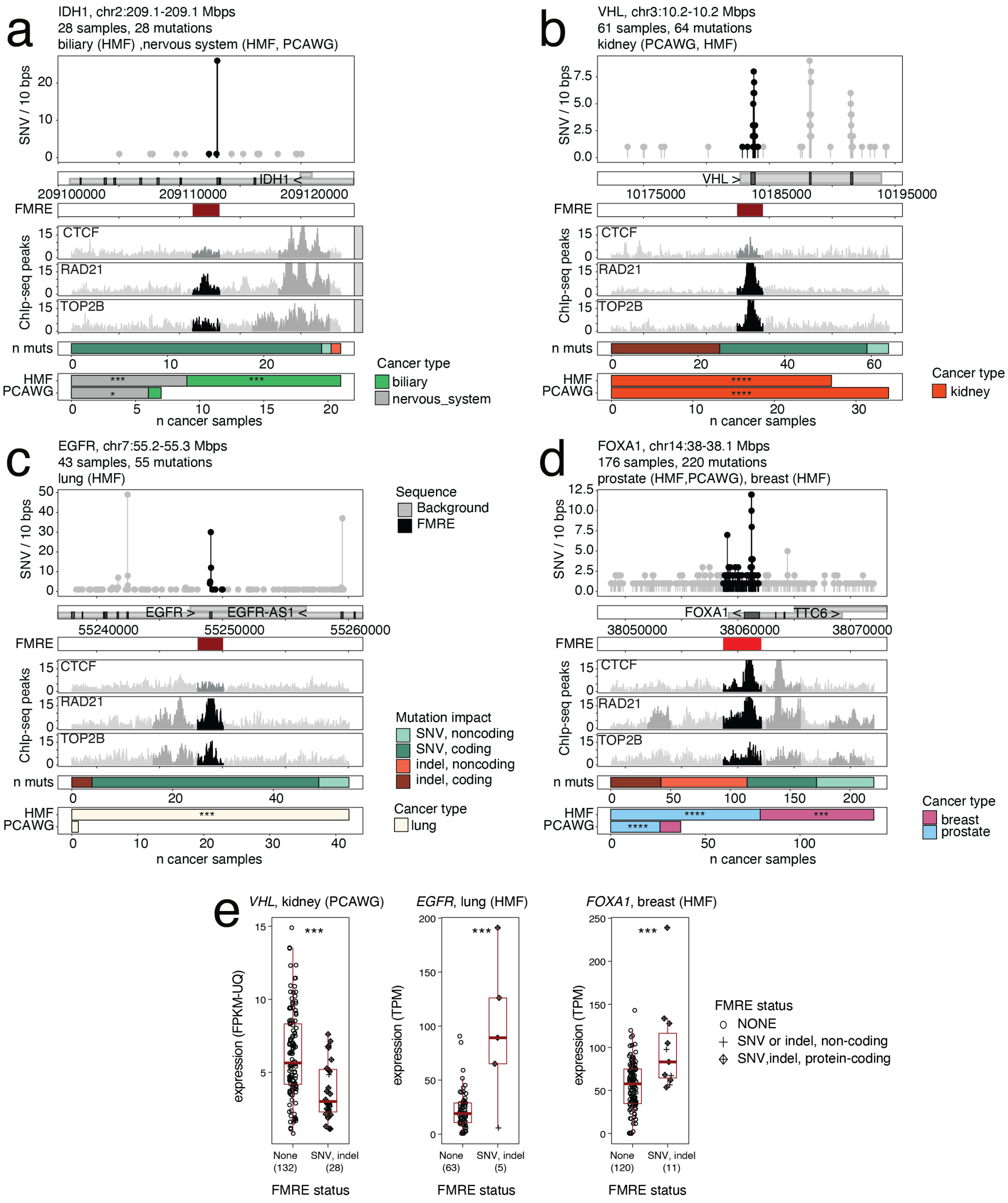


**Figure S5. TOP2B binding sites at hotspot mutations enriched at established cancer genes *EGFR*, *FOXA1*, *IDH1*, and *VHL*. (a-d)** FMREs at hotspot mutations are often bound by TOP2B-CTCF-RAD21 (red) or TOP2B-RAD21 (dark red). Enriched SNVs and indels at the sites are shown (needle plot; top). Representative DNA binding profiles of TOP2B, CTCF, and RAD21 from our ChIP-seq experiments are shown in the middle. Bar plots at the bottom show types of SNVs and indels at the FMRE and the numbers of cancer samples involved. Asterisks at the bar plots show mutational enrichments from ActiveDriverWGS. **(a)** the *IDH1* gene includes an FMRE bound by TOP2B-RAD21 that occurs at the mutational hotspot R132H/C/L that is frequently seen in multiple cancer types. **(b)** The *VHL* gene includes an FMRE bound by TOP2B-RAD21 that is enriched in loss-of-function mutations in primary and metastatic kidney cancer. **(c)** *EGFR* mutations at the hotspots T790M and C797S in metastatic lung cancer. **(d)** In *FOXA1*, the triple site and FMRE located in the final exons and 3'UTR of the gene are enriched in SNVs and indels with protein-coding and non-coding impact in breast and prostate cancer. **(e)** Associations of FMRE mutations and gene expression in matching cancer transcriptomes. Log-transformed expression values were compared using ANOVA analyses and F-tests and accounted for gene CNAs.

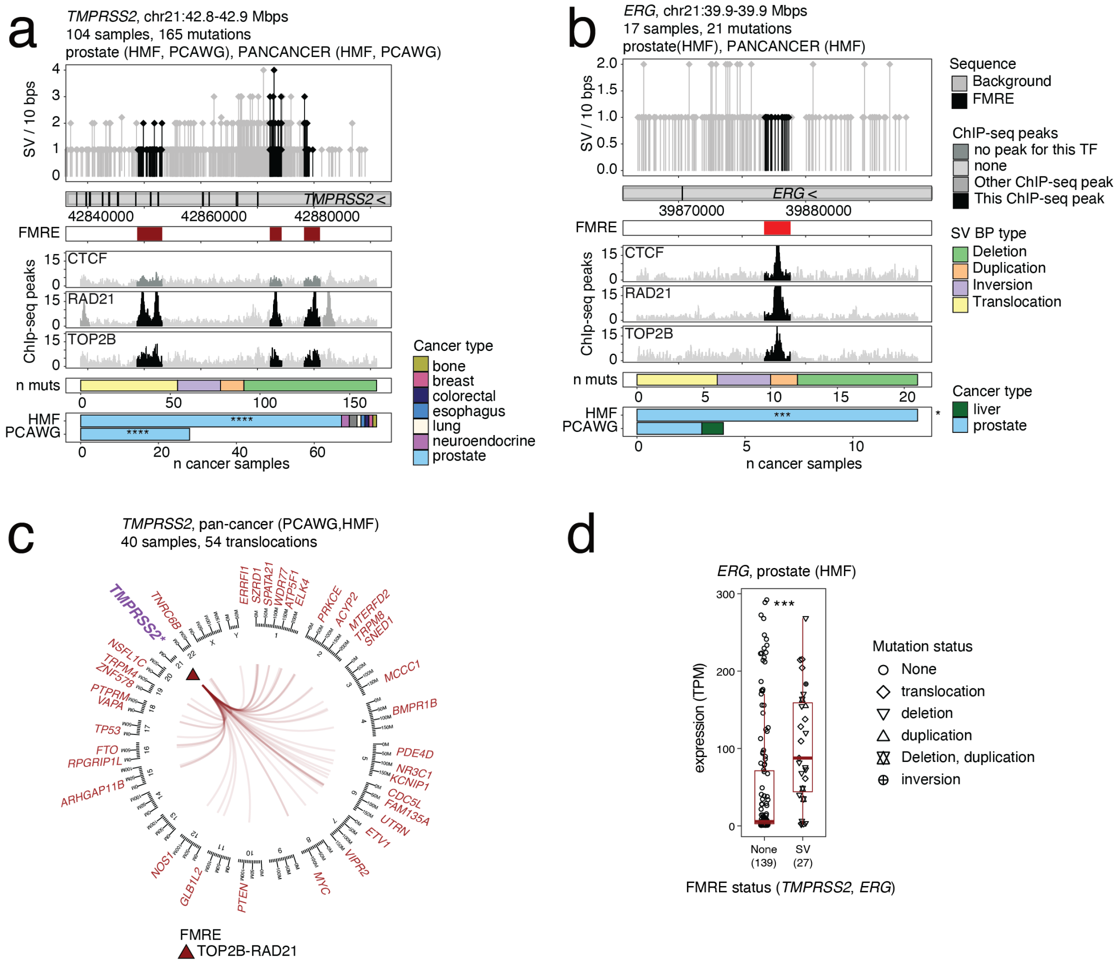

**Figure S6. TOP2B binding site at the *TMPRSS2-ERG* locus is enriched in SVBPs and associates with oncogene activation in prostate cancer. (a-b)** FMREs at *TMPRSS2* and *ERG* SV hotspots are bound by TOP2B-CTCF-RAD21 (red) or TOP2B-RAD21 (dark red). Enriched SV breakpoints at the sites are shown (needle plot; top). Representative DNA binding profiles of TOP2B, CTCF, and RAD21 from our ChIP-seq experiments are shown in the middle. Bar plots at the bottom show types of SVBPs at the FMRE and the numbers of cancer samples involved. Asterisks at the bar plots show mutational enrichments from ActiveDriverWGS. **(c)** Circos plot shows translocations at the TOP2B- RAD21-bound FMRE at the *TMPRSS2* locus with various loci genome-wide. Putative target genes at matching breakpoints are labelled. **(c)** Structural variants at the *TMPRSS2* and *ERG* loci FMRE associate with increased expression of the *ERG* oncogene in metastatic prostate cancer. P-values were computed using log-transformed gene expression values using linear regression and F-tests using matching RNA-seq data from prostate cancer samples in the HMF dataset.

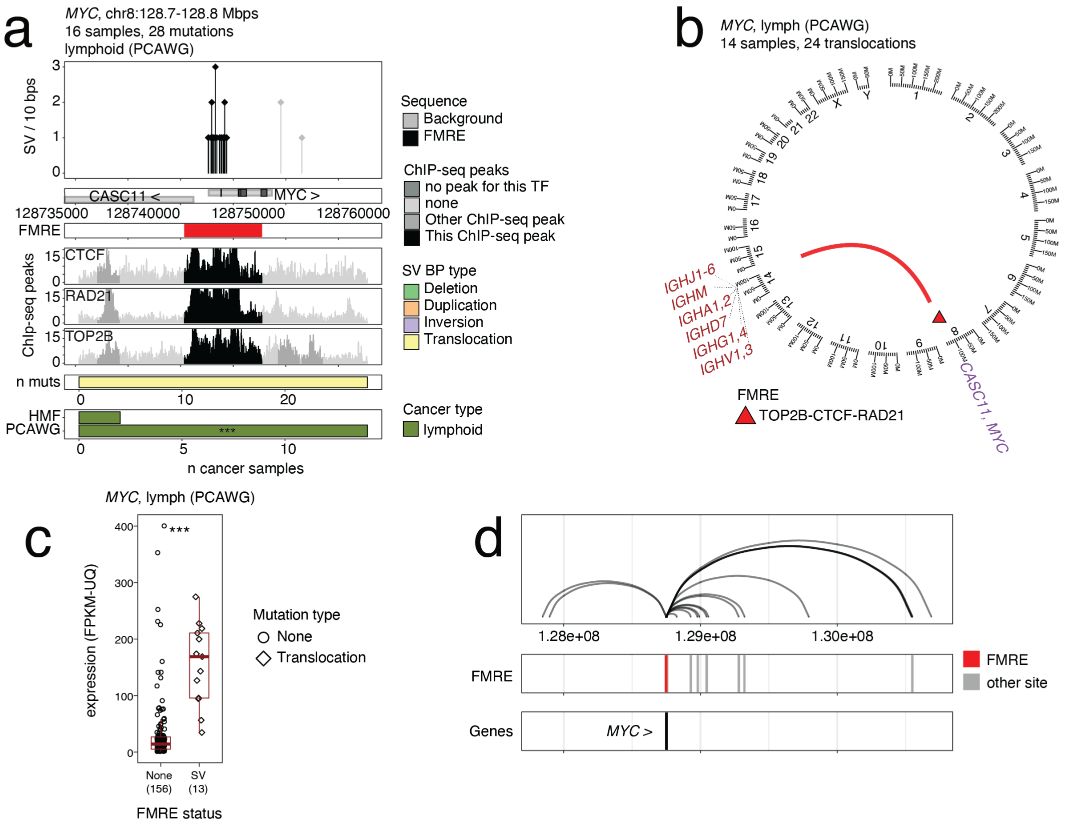


**Figure S7. TOP2B binding site at the *MYC* locus is enriched in SVBPs and associates with oncogene activation in lymphoma. (a)** FMRE at the 3' end of *MYC* is a TOP2B-CTCF-RAD21 binding site (red) with enriched SV breakpoints in lymphoma (needle plot; top). Representative DNA binding profiles of TOP2B, CTCF, and RAD21 from our ChIP-seq experiments are shown. Bar plots at the bottom show types of SVBPs at the locus and numbers of cancer samples involved. Enriched SNVs and indels at the sites are shown (needle plot; top). Asterisks at the bar plots show mutational enrichments from ActiveDriverWGS. **(b)** Circos plot shows recurrent translocations at the TOP2B-CTCF-RAD21-bound FMRE with the immunoglobulin locus on chr14. **(c)** Structural variants at the FMRE associate with increased expression of the MYC oncogene. P-values were computed using log-transformed gene expression values using linear regression and F-tests using matching RNA-seq data from lymphomas in PCAWG. **(d)** The TOP2B-CTCF-RAD21 FMRE at the *MYC* locus shows frequent promoter-enhancer chromatin interactions in multiple types of normal huma tissues. Chromatin loops (top), FMREs and other, non-significant binding sites (middle), and genes with chromatin interactions at transcription start sites (bottom) are shown.

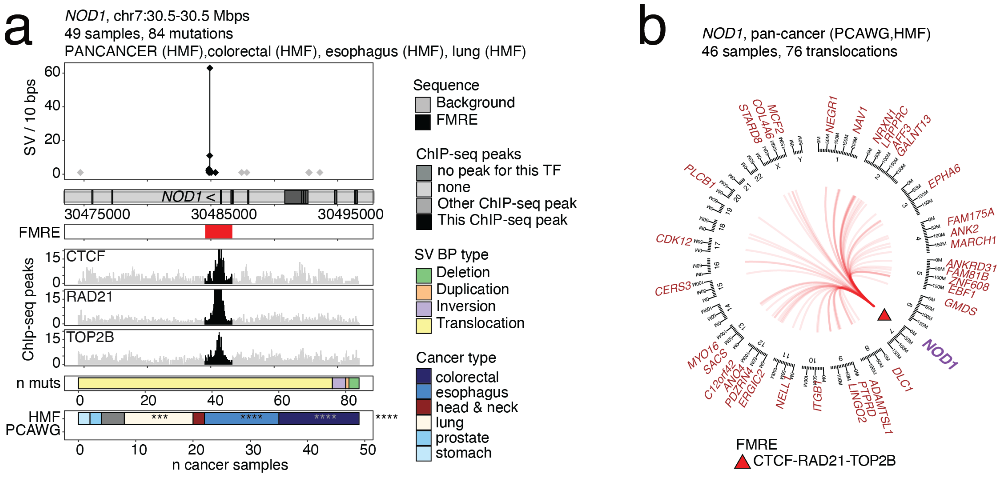


**Figure S8. TOP2B binding site at the putative cancer gene *NOD1* locus is enriched in SVBPs in multiple cancer type. (a)** FMRE at the *NOD1* SV hotspot is bound by TOP2B-CTCF-RAD21 (red). Enriched SV breakpoints at the site are shown (needle plot; top). Representative DNA binding profiles of TOP2B, CTCF, and RAD21 from our ChIP-seq experiments are shown in the middle. Bar plots at the bottom show types of SVBPs at the FMRE and the numbers of cancer samples involved. Asterisks at the bar plots show mutational enrichments from ActiveDriverWGS. **(b)** Circos plot shows translocations at the TOP2B-RAD21-CTCF-bound FMRE at the *NOD1* gene with various loci genome-wide. Putative target genes at matching breakpoints are labelled.

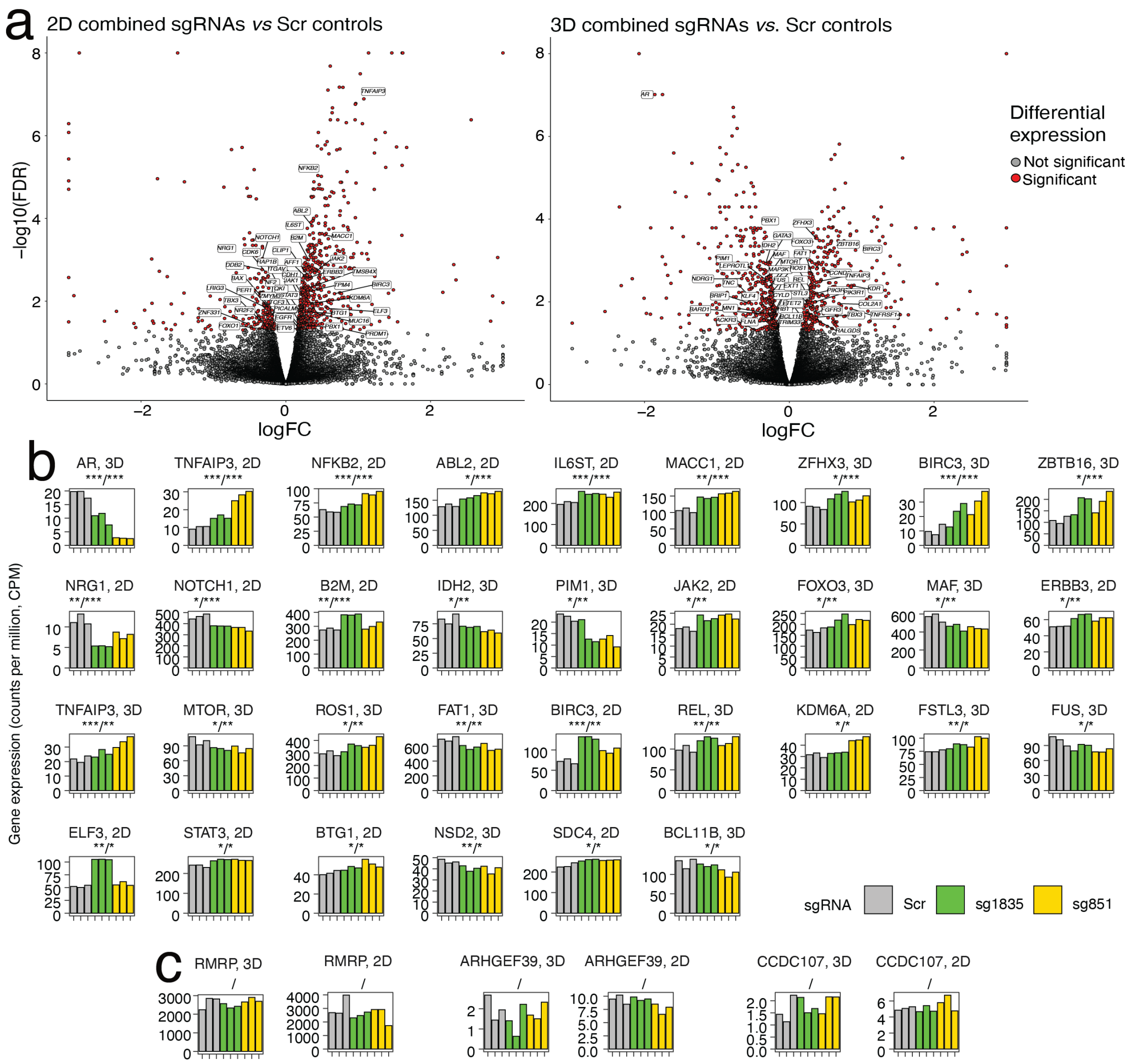


**Figure S9. Transcriptomics analyses of MCF10A cells with FMRE mutations at the *RMRP*/*CCDC107* locus.** **(a)** Differentially expressed genes from RNA-seq profiling of 2D and 3D growth experiments in FMRE-mutant and Scr-treated control cells. Known cancer genes are labelled and adjusted P-values from EdgeR were used to filter genes (FDR < 0.05). **(b)** Known cancer genes from the Cancer Gene Census database with differential expression in either 2D or 3D growth assays shown across the experimental replicates. Asterisks show the FDR value from the assay (left) and the merged FDR value from combining results from 2D and 3D experiments (right). **(c)** The genes RMRP, CCDC107 and ARHGEF39 located at the FMRE locus show no differential gene expression in 2D and 3D growth experiments.
